# Real-Time EEG Neurofeedback as a Tool to Improve Neural Entrainment to Speech

**DOI:** 10.1101/2021.04.19.440176

**Authors:** Francisco Javier Carrera Arias, Nicola Molinaro, Mikel Lizarazu

## Abstract

Neurofeedback represents a particular type of biofeedback whose aim is to teach self-control of brain function by measuring brain activity and presenting a feedback signal in real-time. Traditionally, neurofeedback has been used to complement interventions for various neuropsychological disorders through techniques like frequency training, which attempts to change the power ratio of certain EEG frequency bands. However, to date, there are no neurofeedback approaches that look directly into modulating the neural entrainment to speech. Speech-brain entrainment, which stands for the alignment of the neural activity to the envelope of the speech input, has been shown to be key to speech comprehension. In fact, atypical neural entrainment to speech seems to be consistently found in language development disorders such as dyslexia. Thus, making speech entrainment neurofeedback a promising technique to obtain behavioral improvements. In this work, we present the first open-source brain-computer interface system that can be reliably used to provide speech entrainment neurofeedback while still being flexible enough to deliver more traditional coherence-based neurofeedback. In addition, it has the potential of being an open-source alternative to deliver other types of neurofeedback if configured to do so.

## Introduction

### What is neurofeedback?

Electroencephalography-based neurofeedback (EEG-NF) represents a commonly used technique that involves the real-time processing and measurement of EEG signals while providing feedback to the same person simultaneously (Omejc, Rojc, Battaglini & Marusic, 2018). Through such a feedback loop, that person may attain better control over a neurophysiological parameter of interest by inducing changes in its brain functioning, which may, in turn, affect a given behavior (Omejc et al., 2018).

The origin of EEG-NF goes back to the 1960s were it initially arose a lot of interest due to its potential as a revolutionary clinical intervention (Kamiya, 2011). The technology experienced a decline in use afterwards since it did not meet expectations. However, as technology improved, it saw increasing use again. Today, it is now implemented in a wide array of private clinics over the word (Omejc et al., 2018).

The main function of EEG-NF is to learn how to self-regulate an electrophysiological marker as much as possible. Persons are taught how to increase or inhibit these electrophysiological markers through operant conditioning, which is the process where behavior prevalence is altered through immediate feedback and reinforcement. This premise is rooted on a causal hypothesis stipulating that the deviations in brain function cause the underlying behavioral symptoms leading to the disorders themselves (Sherlin et al., 2011; Sitaram et al., 2016; Enriquez-Geppert, Huster & Herrmann, 2017).

The whole process of EEG-NF works as a feedback loop through a brain-computer interface (BCI), which begins with the acquisition of EEG data from a subject (Figure 1). Immediately after, the EEG signal can be analyzed either offline or in real-time to gather an electrophysiological marker of interest. This marker is then presented back at the subject in visual, tactile, or even auditory form. Traditional examples include video games controlled by the desired brain activity or bar plots displaying the brain activity itself along with a baseline that the subject must try to achieve/improve (Gruzelier, 2014; Enriquez-Geppert et al., 2017).

**Figure 1:**
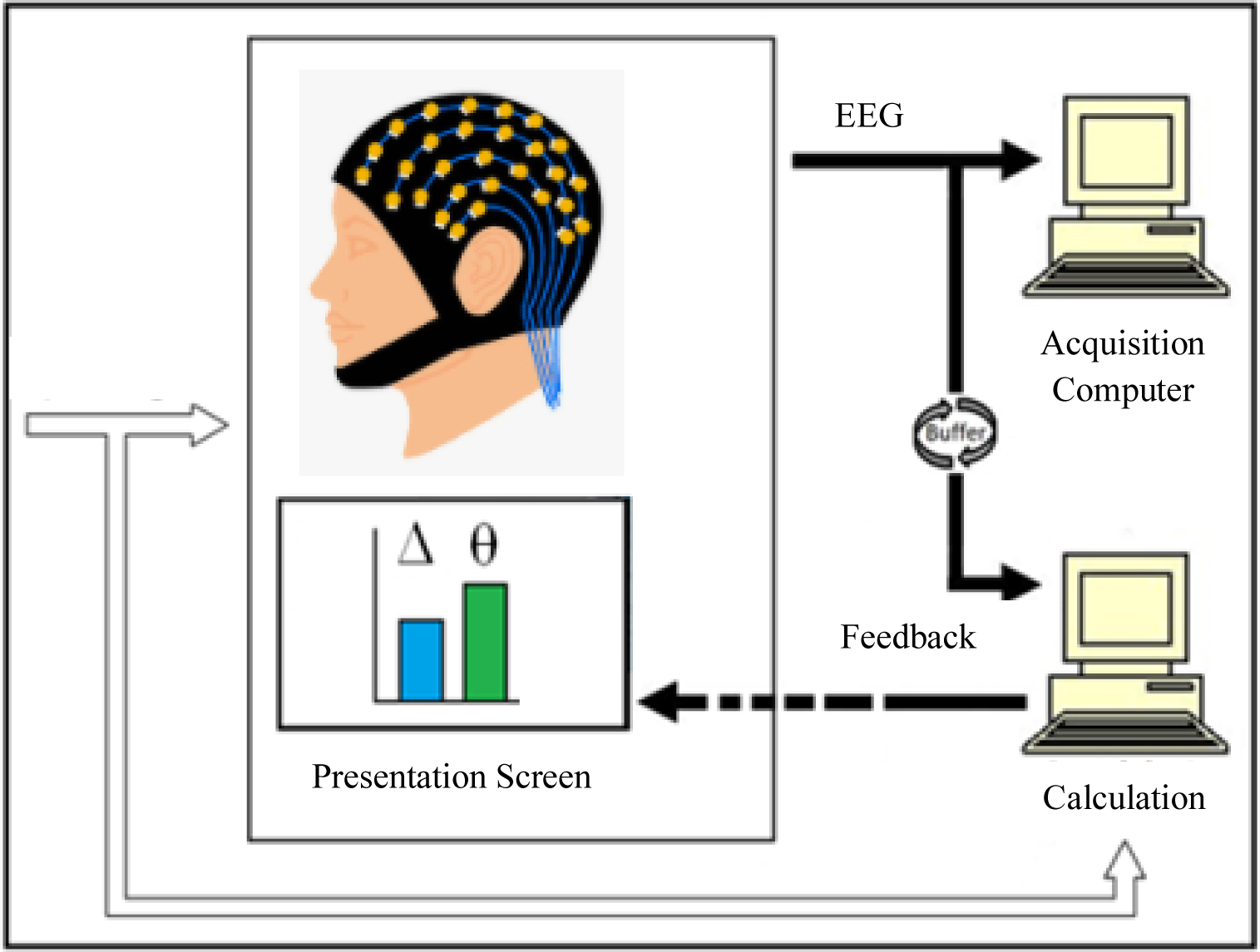
Traditional Neurofeedback BCI Diagram. The Acquisition Computer processes and gathers the EEG data stream. The buffer receives the data from the Acquisition Computer and makes it available to the Calculation Computer. Finally, the Calculation Computer performs the real-time EEG data analyses and presents the metrics back to the participant.

Currently, three major approaches to EEG-NF exist. The one that is by far the most commonly used focuses on frequency training. The aim of this approach is to modulate the power ratio of EEG on-going brain activity, which have been traditionally separated into frequency bands: delta (< 4 Hz), theta (4-7 Hz), alpha (8-13), beta (14-30 Hz) and gamma (> 30 Hz) (Noachtar et al., 1999). The logic behind this kind of neurofeedback modulation is the frequency-to-function mapping. That is, the proposed relationship between the power of certain frequency bands and their associated cognitive functions. The most common frequency training used today is the modulation of the theta/beta ratio that is used for treating attention deficit hyperactivity disorder (ADHD) (Leins et al., 2007).

Alongside frequency training, there are other approaches to EEG-NF. These are primarily the training of slow cortical potentials (SCPs) and coherence-based neurofeedback. SCPs aim to modulate concrete event-related potentials that can be either positive or negative (Birbaumer, 1999). They have been used for patients with epilepsy to increase their seizure threshold levels as well as an intervention for ADHD to improve attentional skills (Birbaumer, 1999; Mayer, Wyckoff & Strehl, 2012). Coherence-based neurofeedback aims to change the connectivity patterns between two or more brain regions defined by a given EEG channel layout (Decker, Fillmore & Roberts, 2017). Distorted connectivity has been shown to appear in several neurologic conditions such as epilepsy, autism spectrum disorder, and traumatic brain injuries, thus leading to the use of this EEG-NF modality in clinical practice (Walker & Kozlowski, 2005; Coben, Wright, Decker & Morgan, 2015; Rostami et al., 2017).

Despite the wide array of clinical applications that EEG-NF modalities possess, there are still gaps not addressed by any of them. One of these, is the fact that the software implementations of most EEG-NF systems are usually not open-source or flexible enough to accommodate more than one EEG-NF modality. They also do not tend to control the temporal delay between neurofeedback presentation with a sufficient level of detail, which is crucial since increased latency significantly affects the sense of agency during BCI control (Evans, Gale, Schurger, Blanke, 2015). More importantly for our purposes however, is that there are no neurofeedback modalities that currently exist to directly modulate speech-brain entrainment.

### A Primer in Speech-Brain Entrainment

Speech-brain entrainment stands for the alignment of neural activity to the slow temporal fluctuations (envelope) of acoustic speech input. The speech signal includes slow temporal fluctuations within the 0.5-10 Hz band that are closely related to phrase and syllable features in the acoustic signal. Tracking such temporal structures, both at phrasal and syllabic rates, is crucial for speech comprehension (Greenberg et al., 2003; Poeppel, 2003; Poeppel, Idsardi, & van Wassenhove, 2008). The phase of low-frequency delta (< 4 Hz) and theta (4 – 7 Hz) oscillations in the auditory cortex synchronizes to the phrasal and syllabic patterns of speech, respectively (Figure 2; Bourguignon et al., 2013, 2020; Molinaro & Lizarazu, 2018; Lizarazu, Lallier & Molinaro, 2019).

**Figure 2:**
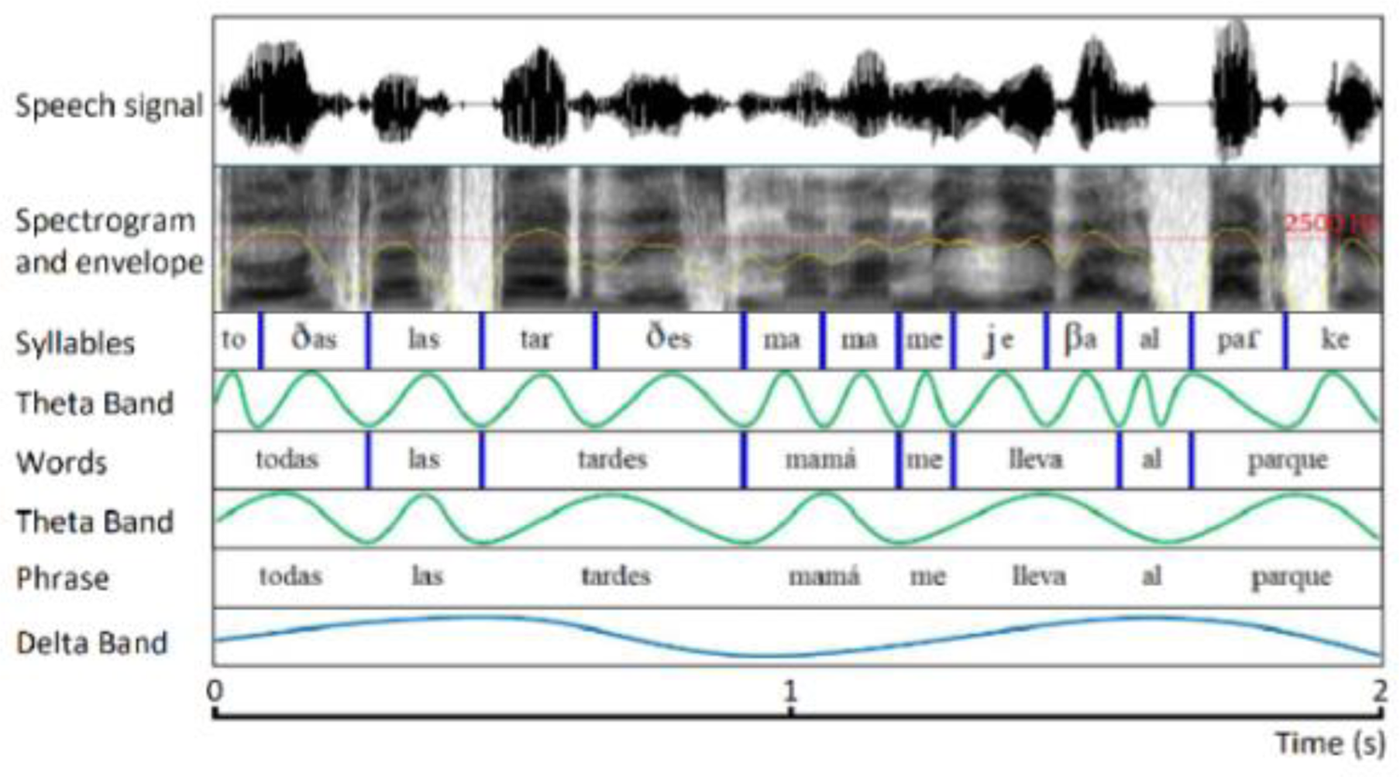
Cortical Tracking of Speech. Representation of the observed synchrony between delta and theta brain oscillations and the phrasal and syllabic patterns of speech respectively (for example: “todas las tardes mama me lleva al parque”/ “every afternoon mom takes me to the park”). On top, we have the original speech wave, followed by its spectrogram containing the envelope shown in yellow. We showcase how theta band oscillations synchronize to syllables and words as well how delta band oscillations entrain to prosodic information.

Speech-brain entrainment reflects to a great extent the bottom-up analysis of the speech signal. Yet, these mechanisms can also reflect top-down modulations of sound processing by using attention, linguistic knowledge, and expectation. Concretely, Park et al. (2015) found that the left inferior frontal and precentral gyri along with posterior temporal areas in the right hemisphere modulate the phase of low-frequency brain oscillations within the left auditory cortex depending on speech intelligibility. Moreover, they noted that speech-brain entrainment is improved as a function of stronger top-down modulation. Combined, these results ultimately suggest that theta and delta brain oscillations are crucial in exerting top-down control during speech processing. These findings are further supported by a plethora of other studies that have also suggested that neural entrainment to speech correlates with the intelligibility of the speech signal (Ahissar et al., 2001; Ding & Simon, 2013; Doelling, Arnal, Ghitza & Poeppel, 2014; Gross et al., 2013; Peelle, Gross, & Davis, 2013; Pérez et al., 2015; Rimmele, Golumbic, Schröger, & Poeppel, 2015; Ghinst et al., 2016; Molinaro & Lizarazu, 2018; Lizarazu et al., 2021; Molinaro et al., 2021). For instance, in Ghinst et al (2016), participants were asked to pay attention to a speech stream embedded within a “cocktail party” multitalker background noise at different intensity levels. They found that delta and theta neural entrainment was stronger for the attended speech compared to the multitalker background. In addition, they showed that neural entrainment to the attended speech decreased as the level the multitalker background noise progressively increased.

### Clinical Neurofeedback Applications of Speech-Brain Entrainment

Atypical entrainment to speech has been shown to play a role on various neuropsychological disorders such as dyslexia, stroke, and Broca’s aphasia (Goswami, 2011; Molinaro et al., 2016; Feenaughty et al., 2017; Liberto et al., 2018; Thors, 2019; Lizarazu et al., 2021b,c). For example, in terms of developmental dyslexia, Goswami (2011) theorized that atypical speech-brain entrainment could be part of the underlying cause. She stipulated that impairments in the temporal sampling of speech by brain oscillations could explain the phonological and perceptual difficulties that characterize individuals with dyslexia.

In this vein, numerous functional brain studies have shown atypical neural entrainment to speech in dyslexia (Hämäläinen, Salminen, & Leppänen, 2013; Lizarazu et al., 2015; Cutini et al., 2016; Molinaro et al., 2016; see Jiménez-Bravo, Marrero, & Benítez-Burraco, 2017 for a recent review). Molinaro and colleagues (2016) found that dyslexic readers had impaired neural entrainment in the lower delta band range localized in both the left inferior frontal gyrus and right auditory cortex. Moreover, they found impaired functional connectivity between oscillations in the left frontal cortex and the right auditory cortex. Lizarazu and colleagues (2015) further reported atypical neural entrainment to non-linguistic auditory signals (amplitude modulated white-noise) in the theta band in dyslexics. Together, these results suggested that individuals with developmental dyslexia display reduced neural entrainment to low-frequency speech oscillations compared to controls while listening to speech.

This could be a clear example where speech entrainment neurofeedback could have a huge impact by training the ability of individuals to better synchronize to a given speech signal. Specifically, neurofeedback may be able to enhance the top-down modulation of speech-brain entrainment in auditory regions (Park et al., 2015). Therefore, to that end, we have developed a first of its kind open-source BCI toolbox in Matlab that is capable of providing real-time speech entrainment neurofeedback while participants listen to an audio signal. This required a very precise design capable of aligning EEG data to the exact segments of audio that the participant is listening to while performing all the neurofeedback calculations in real-time. Furthermore, we built the toolbox such that it is flexible enough to accommodate more traditional coherence-based neurofeedback with respect to any given EEG channel, as well as to provide other types of neurofeedback if configured to do so.

In the following sections, we will describe how this toolbox has been implemented and how it operates in detail under the simulated data framework we used to test it. Moreover, we also propose a speech-in-noise experiment to test its implementation using real data, which is crucial for any preliminary tests on clinical populations.

## Methods

Overall, we conceptualized a complete neurofeedback session using our toolbox as a two-step process. The first step involves an offline localizer experiment that aims to find the EEG sensors showing the highest levels of speech entrainment. The second step is the neurofeedback session itself, which consists of continuously presenting the strength of the neural entrainment to speech. The objective is to maximize the neural entrainment to speech across the neurofeedback session. In the following sections, we will go into the detail of each of the steps showing how the toolbox works internally, as well as to provide a guide on how to conduct successful neurofeedback sessions. All the code developed for this toolbox can be seen in Appendix A as well as in the following URL to Matlab File Exchange: https://es.mathworks.com/matlabcentral/fileexchange/78170-real-time-speech-brain-entrainment-neurofeedback-toolbox?s_tid=LandingPageTabfx

### EEG Data Acquisition

This toolbox has been designed to work with BrainAmp amplifiers and BrainVision Recorder software (BrainProducts, Germany). Based on our testing setup, we recommend using EEG systems with 32 electrodes positioned according to the international 10-20 system (Jasper, 1958). Moreover, we recommend placing the reference channels in the earlobes and keeping scalp-electrode impedance below 8 kΩ to ensure high-quality EEG recordings. The sampling rate of the EEG system should be at or above 500 Hz.

### Part 1: Offline Localizer Session

The localizer part of the neurofeedback session behaves very much like a classical offline EEG experiment. The objective, as mentioned previously, is to locate the EEG sensors that display the highest level of neural entrainment with the speech envelope for a given frequency band. This step is optional, but highly recommended since it is customary for EEG-NF applications to focus on a few channels of interest (Omejc et al., 2018). By default, the frequency range of interest is between 0.5-7 Hz, which corresponds to the delta and theta EEG bands. This default was chosen given the extensive literature highlighting their importance in speech-brain entrainment (Bourguignon et al., 2013; Molinaro et al., 2016; Molinaro and Lizarazu, 2018; Lizarazu, Lallier and Molinaro, 2019).

To run the localizer session, we developed the function *speech_brain_coherence.m,* which is built on top of Fieldtrip (Oostenveld, Fries, Maris & Schoffelen, 2011), and it is able to perform all the necessary steps. These include:

1. Preprocessing and trial segmentation of the offline EEG experiment data.
2. Calculation of the degree of neural entrainment based on the coherence between each EEG channel and the envelope of the speech presented across trials.
3. Gathering the *n* EEG channels with the highest average coherence between the range of frequencies of interest defined by the user.
4. Optionally, as an additional step, the user can choose to plot the coherence across the range of frequencies of interest. An example of this can be seen in Figure 3.

**Figure 3:**
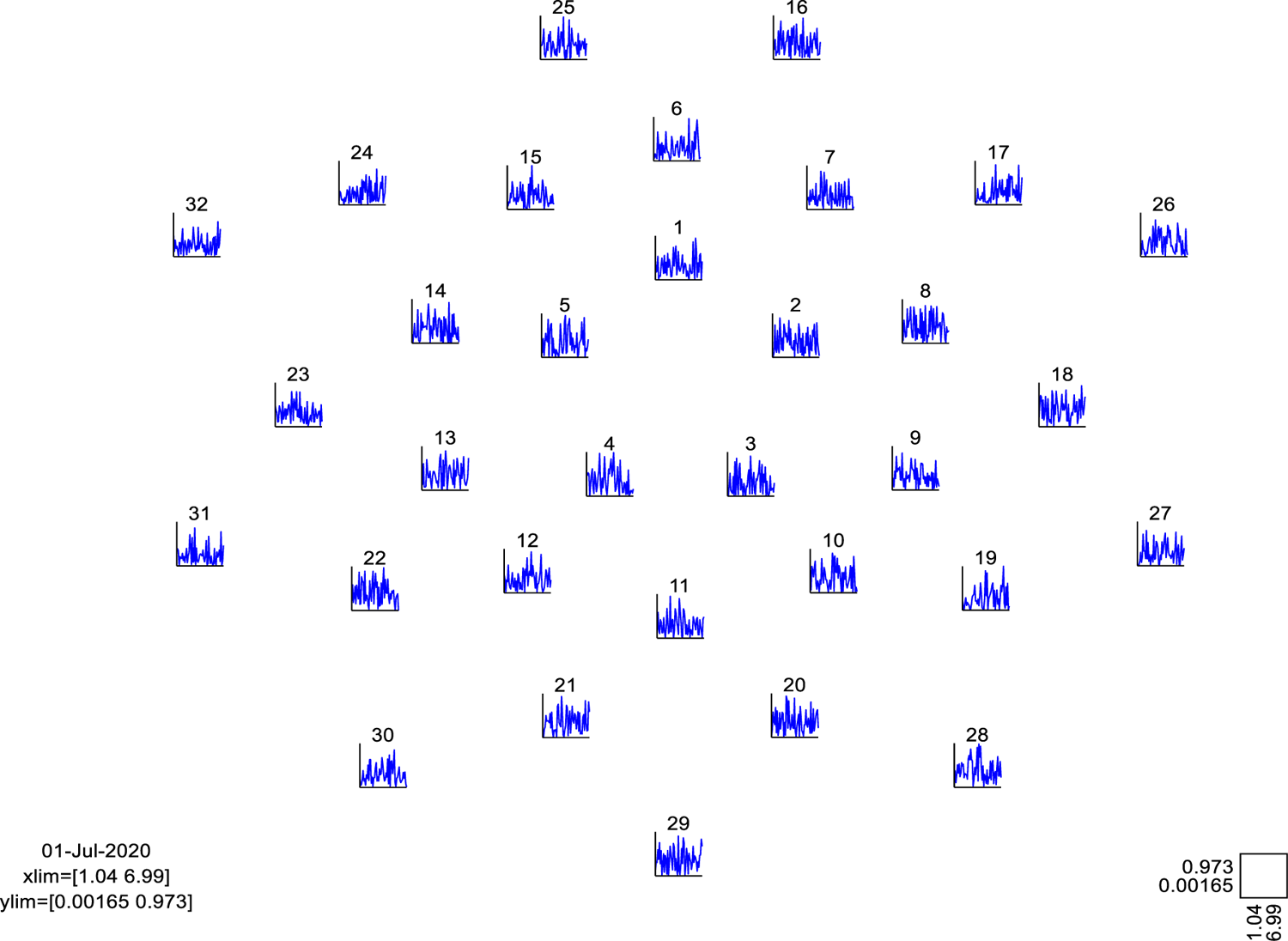
Localizer session. In this case, the range of frequencies of interest is from 0.5-7 Hz, which correspond to the delta and theta EEG bands. Results are shown using a 32-channel EasycapM7 layout. The coherence shown represents that obtained from calculating the coherence between simulated EEG data and the speech envelope.

The current implementation of the toolbox only segments the trials based on the trigger values and applies a FIR band-pass filter to the data using the frequency band of interest given by the user. Ideally, this frequency band will correspond to the frequencies tracked during the neurofeedback session. The *Trial_Segment.m,* function, which is in charge of this step, is very flexible and can easily be modified to accommodate more preprocessing steps like re-referencing as well as electrooculogram or motion artifact detection if desired.

### Coherence

For this session, we used coherence to quantify the degree of neural entrainment to speech. Coherence measures the phase synchronization between two signals (i.e. speech envelope and EEG data) in the frequency domain. This measure was chosen for the localizer since it provides a fine-grained view of entrainment at each individual frequency within the band of interest. This can be useful to pick sensors of interest as well as for standalone offline data analyses. The definition of coherence between two signals *x(t) and y(t)* can be seen in Equation 1:

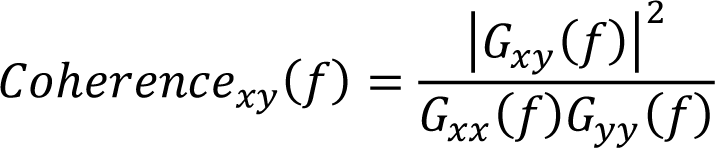

Where *f* stands for frequency in Hz, *G_xy_*(*f*) stands for the cross-spectral density between the signal of the EEG channel and the speech envelope, *G_xx_*(*f*) and *G_yy_*(*f*) correspond to the spectral density of the EEG channels and the speech envelope with themselves, respectively (Bruña, Maestú & Pereda, 2018). For obtaining the speech envelope, we obtained the magnitude of the analytic signal of the speech after a Hilbert transformation. Following that, we downsampled the signal to the sampling frequency of the given EEG system, which in our case was 500 Hz. Coherence values range from 0 to 1 where 0 indicates no phase synchrony and 1 indicates perfect phase synchrony.

It is important to highlight that the user can select the *n* number of channels to keep for use during the neurofeedback session. As a guideline, we suggest selecting 3-5 sensors of interest at the most. It is likely that only a few would be needed given the custom in neurofeedback applications (Omejc et al., 2018). As an additional optional feature, the user can also generate interactive plots of the coherence spectrum for each channel (Figure 3). This allows for a thorough study of a participant’s speech entrainment prior to any neurofeedback session if desired. Thus, even though the *speech_brain_coherence.m* function has been designed primarily with speech-brain entrainment neurofeedback in mind, it can also work as a reusable standalone function for offline experiments.

### Part 2: Real-Time Speech Entrainment Neurofeedback Session

The bulk of the work conducted has been related to the implementation of a brain-computer interface (BCI) system that could make reliably speech entrainment neurofeedback with minimal delay. To that end, we developed a series of functions built on top of several toolboxes. These being Fieldtrip (Oostenveld et al., 2011), Psychtoolbox-3 (Brainard, 1997; Pelli, 1997; Kleiner, Brainard & Pelli, 2007) and a slightly modified version of the TCP/UDP/IP toolbox for Matlab (Rydesäter, 2019).

### BCI Data Pipeline Setup

For running a successful neurofeedback session, the BCI data pipeline depicted in Figure 4 must be in place. The envisioned data pipeline begins with a real-time EEG data stream from a BrainAmp amplifier, which is recorded in the acquisition computer running BrainVision Recorder (BrainProducts, Germany). Prior to the neurofeedback session, the remote data access (RDA) server within BrainVision recorder’s preferences must be activated (Figure 5). This allows the EEG data to be available through a TCP/IP connection to the data processing and presentation computer.

**Figure 4:**
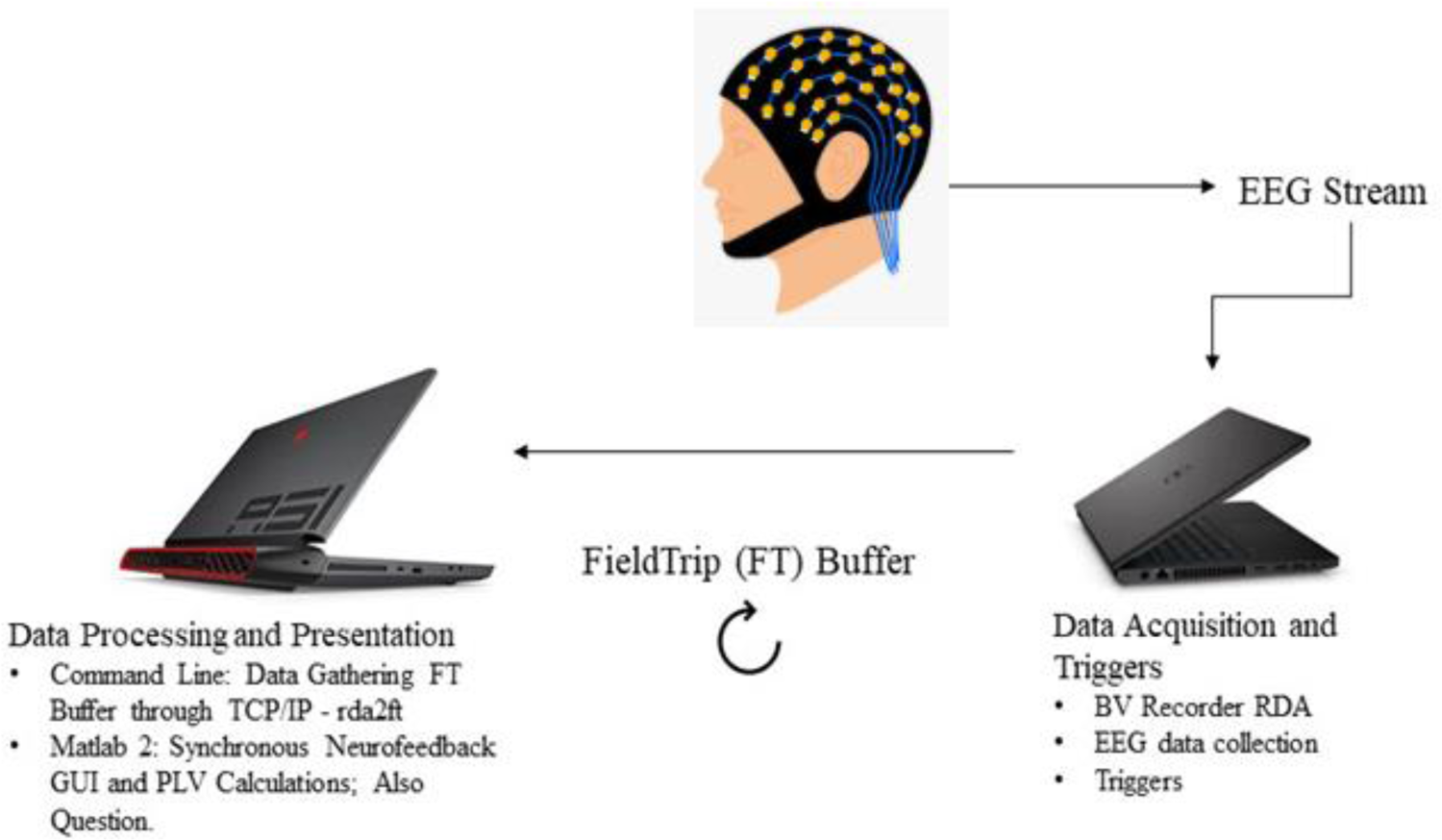
BCI setup in the present study. BV stands for BrainVision, RDA stands for remote data access, TCP/IP stands for transmission control protocol and internet protocol. GUI stands for graphical user interface.

**Figure 5:**
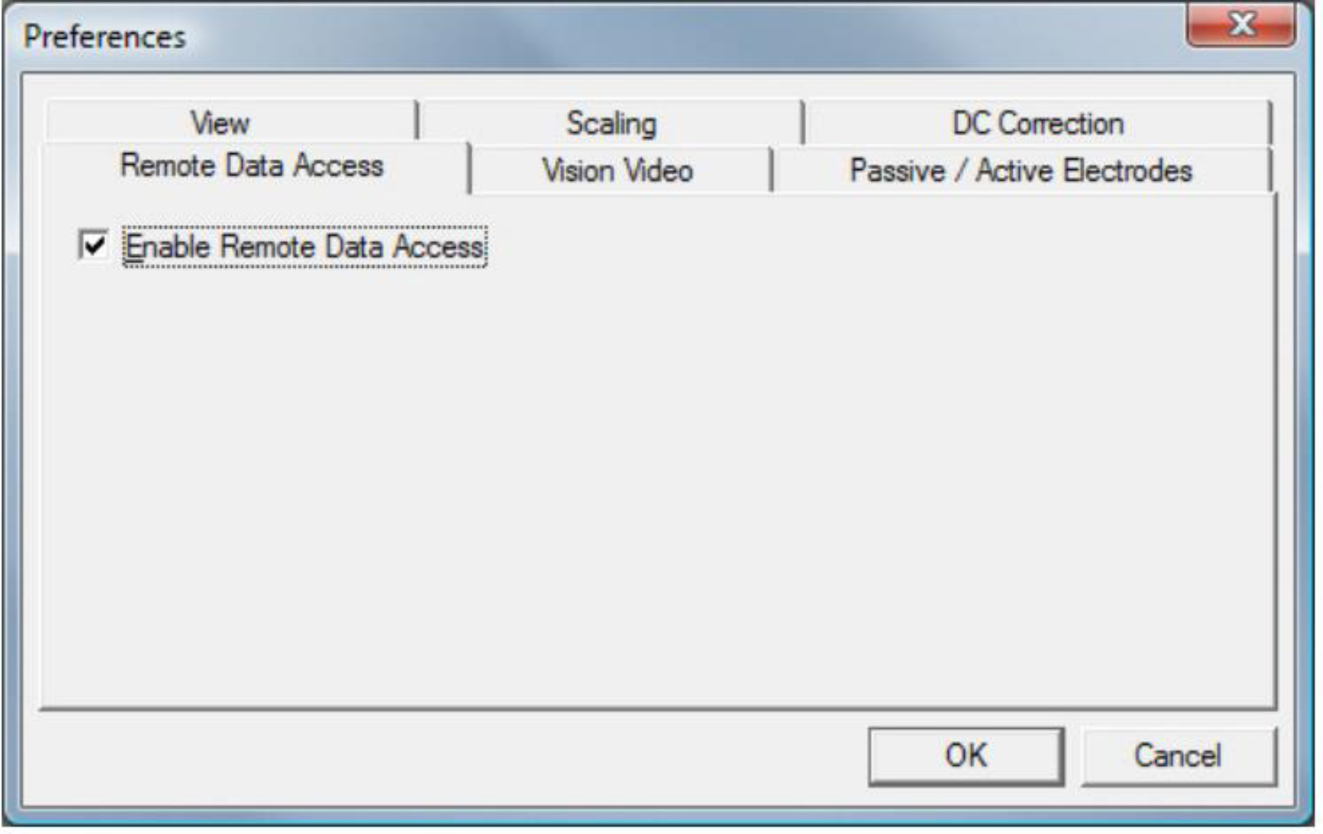
Location of the RDA activation checkbox with BrainVision Recorder’s Preferences.

The acquisition computer will also house the Matlab script contained within this toolbox to send the trigger pulses (*Trigger_Gen_Multi.m*). This will allow the data processing and presentation computer to synchronize the neurofeedback with the speech/audio stimuli. The current toolbox implementation uses a mex-file parallel port interface to send the triggers to the EEG system (Schieber, 2020). However, this can be a researcher’s choice and other options could be used instead. The only requirement is that the software is able to send TTL pulses to an EEG system.

Moving on from the acquisition computer, we have the utilities that are housed within the data processing and presentation computer. The key utility that makes the BCI data pipeline implemented in this toolbox possible is Fieldtrip’s data buffer. The buffer houses the data streamed from the RDA server in BrainVision Recorder. To collect the data from the RDA server, we used a compiled C utility that is provided by Fieldtrip called *rda2ft* (Oostenveld et al., 2011). In order to activate *rda2ft*, the IP of the acquisition computer is needed. Thus, it is key that the data processing computer and acquisition computer are in the same network. The port of the RDA server is also required, which for a x64 bit architecture computer is port 51244. By default, once given those parameters, *rda2ft* instantiates the Fieldtrip buffer on the local data processing and presentation computer on port 1970. Being a C-built utility, *rda2ft* is extremely fast, and it is capable of collecting data from the RDA server every 2 ms. This allows the stimulus presentation and subsequent neurofeedback processes to begin immediately after the trigger is sent within the acquisition computer.

### Phase-Locking Value

The aim of the neurofeedback session is to continuously present the strength of the neural entrainment to the speech, which during the neurofeedback session is quantified using phase-locking value (PLV). Specifically, we designed the toolbox to compute the average PLV between brain oscillations at the delta (0.5-4 Hz) and theta (4-7 Hz) frequency bands and the speech envelope.

Coherence was preferred for the localizer session since it provides a more detailed view of entrainment within each individual frequency inside a band. This can give a researcher more control in terms of selecting the EEG sensors of interest and standalone value as a reusable function for offline experiments. However, for the neurofeedback session we chose PLV because of the opposite. As implemented, it outputs a single value for an entire frequency band, which is desirable for the delta/theta average calculation we use for neurofeedback. It is also a slightly leaner calculation, which helps in terms of computation speed (Bruña et al., 2018). Since the neurofeedback session calculates the PLV in real-time, (2) it was defined as shown in Equation 2 for the purposes of our toolbox:

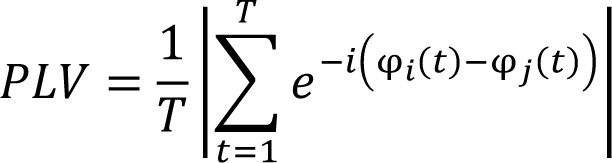

Where *T* is the data length (number of samples), φ_i_(t) is the instantaneous phase of the signal of each EEG channel at time *t*, and φ_j_(t) is the instantaneous phase of the speech envelope (default) or any reference channel of choice at time *t*. The instantaneous phases of each of the signals and the speech envelope were obtained from the phase angle of the analytic signals after a Hilbert transformation and a FIR band-pass filter.

PLV is defined as the absolute value of the mean phase difference between two signals, and has values ranging from 0 indicating no phase alignment to 1 indicating complete phase alignment (Lachaux, Rodriguez, Martinerie & Varela, 1999; Bruña et al., 2018). The goal of a participant during the neurofeedback session, by default, will be to maximize as much as possible the theta/delta PLV average across the channels of interest gathered from the localizer over time.

### Experimental Design and Neurofeedback BCI Loop

Once the data is being collected into the buffer through *rda2ft*, the heavy lifting of presenting the neurofeedback along with the speech/audio stimuli falls to the *ft_realtime_plv_fully_sync.m* function. As a general overview, *ft_realtime_plv_fully_sync.m* constantly checks the buffer for events (triggers) of interest selected by the researcher. While no triggers of interest are detected, a white fixation cross is displayed on the screen. During this time, participants can be invited to blink to reduce artifacts during the neurofeedback presentation since these data are not analyzed. Once a trigger of interest is detected, it proceeds to play a piece of audio that is randomly selected among the ones contained within the stimuli folder associated with the trigger value (see the documentation for *Audio_Processor.m* for further details into how stimuli folders must be named). The real-time neurofeedback itself will begin once there is enough EEG data in the buffer associated with the presentation of the speech/audio. It is important to note that the determination of whether there is enough EEG data is controlled by the chosen presentation frequency window. This is a key user-defined parameter that defines how often the neurofeedback should be presented.

Given that the speech/audio is an external signal to the EEG, we were extremely careful to align each segment of real-time EEG data to the exact segment of the speech/audio that an individual listened to during data collection. As such, once an event of interest is detected, the *ft_realtime_plv_fully_sync.m* function calls *ft_trialfun_speechwindow.m* (a custom trial definition function) to define the boundaries of the neurofeedback presentation windows. We use the EEG data stream samples from the buffer to mark the beginning and the end of these presentation windows. Likewise, these boundaries are used to gather the exact segment of the speech/audio envelope that corresponds to what the individual listened to during that time in order to perform accurate PLV calculations.

It is crucial to note that the *ft_trialfun_speechwindow.m* function will only create windows that are exact divisions with respect to the audio length. That is, if we have an EEG system with a sampling rate of 500 Hz, speech stimuli lasting 10s, and a window size of 2s, the *ft_trialfun_speechwindow.m* will create 5 trials. All of these would contain 1000 samples and will cover the entirety of the audio envelope. However, if we have speech stimuli lasting 11s, the ft*_trialfun_speechwindow.m* will also create 5 trials containing 1000 samples each since 10 is the nearest exact division of 2. This is done to avoid confounding effects of window length by guaranteeing that the presentation windows are always consistent.

During the presentation of the neurofeedback, it is important to emphasize the global moving average baseline. This baseline is plotted as a horizontal line in the bar chart that is used as the default neurofeedback visualization method. We included this baseline in order to provide a clear improvement objective to the individual receiving the neurofeedback. The default instruction is to try to move that line as closer to 1 (perfect phase alignment) as possible.

Once the last presentation window is reached, the neurofeedback will remain in the screen for the duration of another presentation window. This way, the participant receiving the neurofeedback will have enough time to process the information presented in the last window. After this, the *ft_realtime_plv_fully_sync.m* will allow researchers to evaluate participant behavior. By default, the toolbox comes with an intelligibility question in which the participant has to evaluate in a scale from 1-9 how intelligible they found a given speech/audio stimulus. However, other options are possible such as yes/no comprehension questions. In any case, the function waits for a keyboard key press as a response before returning to the white fixation cross. This fixation cross will remain in the screen until the next trigger of interest is detected and the cycle is repeated again.

The BCI loop will continue until all the stimuli have been presented. Once that point is reached, the BCI loop ends and the function returns the results of the neurofeedback session. These results are stored in a Matlab cell array format with as many rows as stimuli and columns containing the condition based on the trigger value, the answer to any behavioral questions, and the average theta/delta PLV at each presentation window. A visual representation of the BCI loop explained above can be seen in Figure 6.

**Figure 6:**
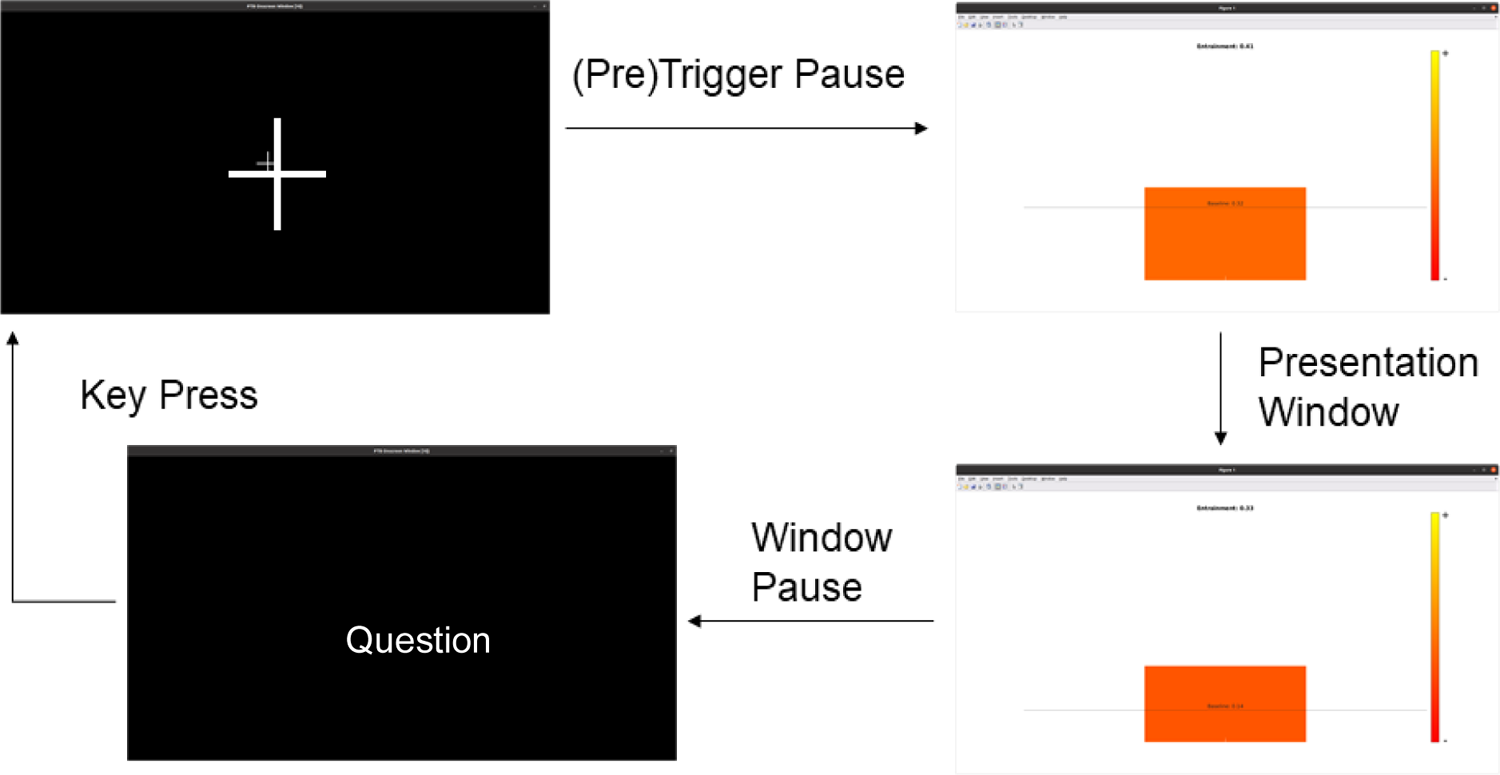
BCI presentation diagram. The (pre) trigger pauses stand for the time in between triggers, which is a parameter controlled by *Trigger_Gen_Multi.m* in the acquisition computer. This pause serves to control the presentation of stimuli and provide time for participants to blink. The presentation windows, as mentioned, are how often the neurofeedback presentation is desired (i.e every 2s); the window pause is also the same amount. Lastly, behavioral questions (for example, a comprehension question) are controlled by key presses. As soon as the participant presses a key, the system goes back to fixation.

### Neurofeedback Computational Performance

Our testing was run on an Alienware Area 51m laptop as the data processing and presentation computer. This computer contained an Ubuntu 20.04 operating system, a 9^th^ generation Intel i9-9900k CPU, a Nvidia RTX 2060 GPU, 32 GB of RAM and a 17.3 inch screen with a 144 Hz refresh rate. Under these specifications, we obtained an overall average delay of 218.9 ms for all the signal processing and PLV calculations, which is a good real-time performance for a Matlab toolbox.

We obtained this value by testing across 3 different presentation windows, which we hypothesize to be the most likely to be used, and 5 channels of interest from the localizer. We do not expect that a researcher would use that many channels given that neurofeedback applications tend to focus on less (Omejc et al., 2018), but it allowed us to see how the toolbox would perform under harder circumstances. Figure 7 shows the details of the calculation timings for each of the presentation windows tested.

**Figure 7:**
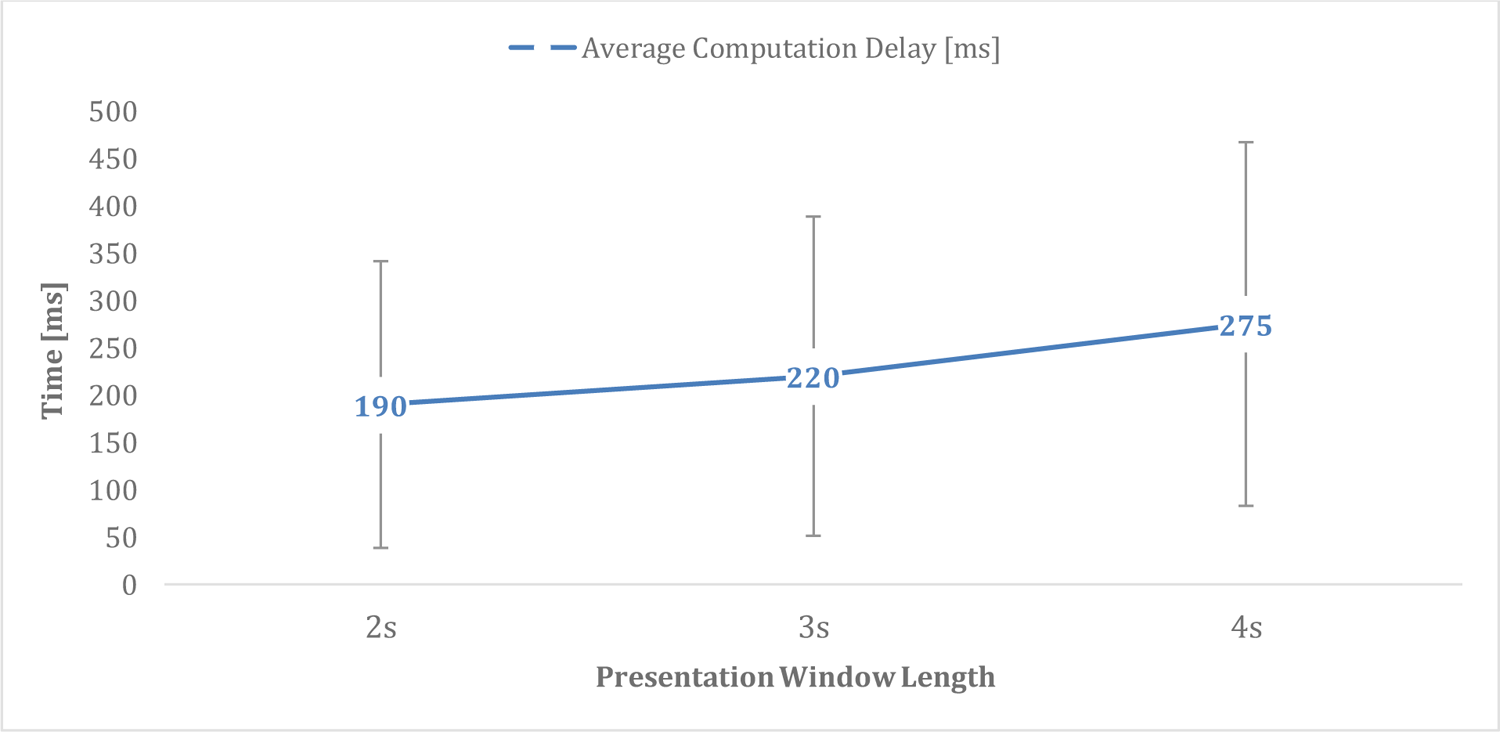
Computation Delays. Average computation delays in milliseconds by presentation window length. As can be expected, the longer the presentation window, the longer the average delay. Error bars are standard deviation representing the variability in computation times. These can be caused due to ongoing background processes within an operating system and Matlab’s optimizations after the first time a given piece of code is executed.

### Proposed Validation Experiment

Due to technical difficulties that arose during the time this work was produced (closure of the labs to prevent the spread of the virus COVID-19), we could only test and validate the operation of our neurofeedback toolbox using simulated EEG data. However, in order to better position speech-brain entrainment neurofeedback as a strong candidate for clinical application, a formal experiment with real data is needed.

In order to achieve this and provide a stronger rationale for using our neurofeedback approach with clinical populations in the future, we first propose a speech-in-noise experiment with healthy participants. This study would contain two groups. A control group, which would receive random neurofeedback, and an experimental group that would receive the true average theta/delta PLV neurofeedback as they listen to speech. Given the sample sizes of prior entrainment experiments, each group would contain 10 participants (Ghinst et al., 2016). Thus, resulting in a total sample size of 20 healthy subjects.

The experiment would consist of 4 speech-in-noise conditions that are randomly presented during the neurofeedback session. Clean speech would be presented during the localizer session. The stimuli would be 20 seconds long with neurofeedback presented in windows of 2s. After this presentation, participants would be asked to rate the intelligibility of the speech as well as comprehension questions to gather behavioral measures. It is important to note that individuals would be asked to attend to the voice belonging to the person whose clean speech was listened during the offline localizer session while in the neurofeedback session (Ghinst et al., 2016). This would be done in order to provide a clear attention target for top-down entrainment modulation. The noise would be a cocktail party noise of 6 people (3 females). A full diagram of the conditions that could be used for this experiment can be found in Figure 8.

**Figure 8:**
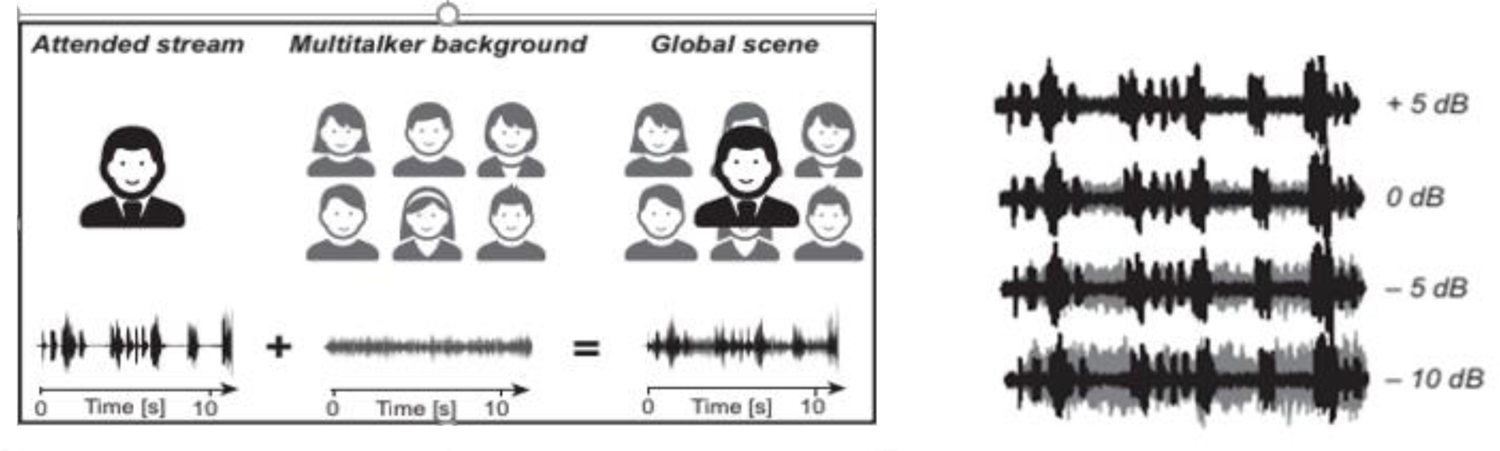
Proposed Speech-in-noise conditions for neurofeedback validation in healthy participants. Adapted from Ghinst et al., 2016.

If neurofeedback can indeed modulate top-down speech-brain entrainment in our proposed experiment, we would expect results similar to those shown in Figure 9. That is, we would expect to observe strong positive correlations at the individual level between the mean delta/theta PLV and the intelligibility scores of each stimulus presented during mild noise conditions on participants receiving true neurofeedback. We also hypothesize that the participants in the control group will display weak correlations between their intelligibility scores and the mean delta/theta PLV across presentation windows. Furthermore, we expect neurofeedback to decrease in effectiveness as background noise increases given the decrease in entrainment observed in past studies (Ghinst et al., 2016).

**Figure 9:**
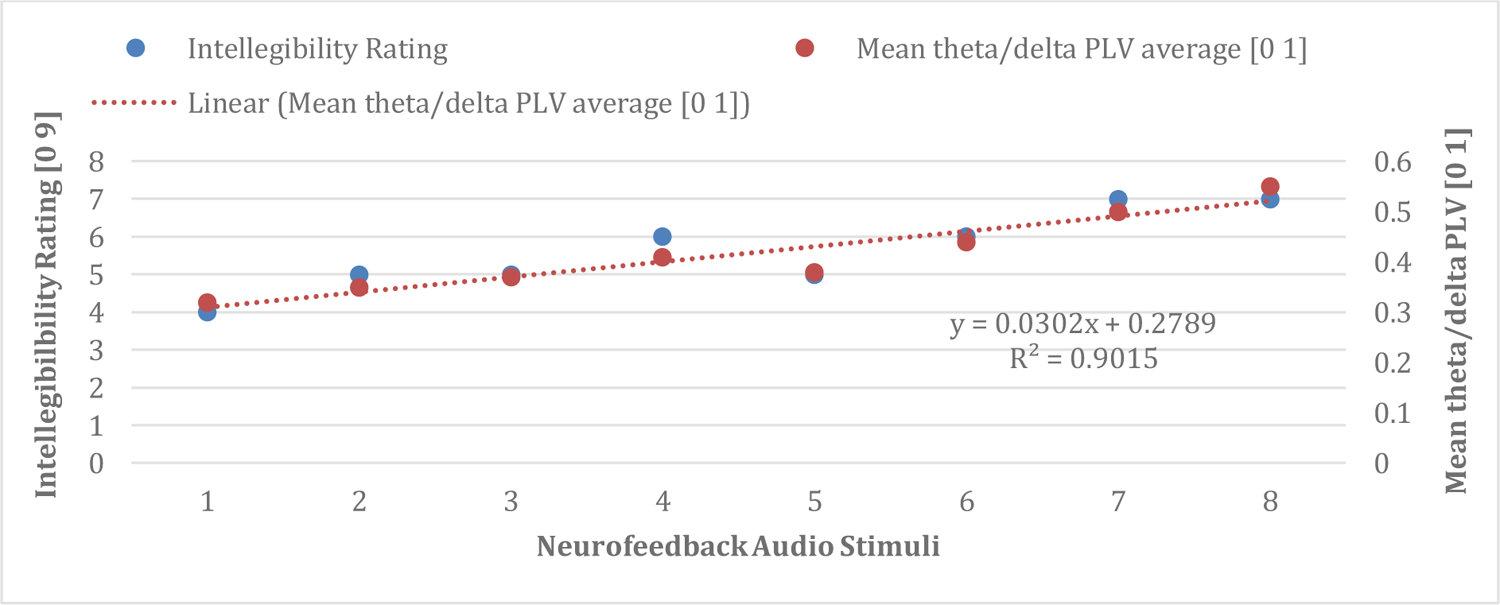
Hypothesized effect of speech-entrainment neurofeedback on stimulus intelligibility over time in a single individual during a mild noise condition such as +5 dB.

## Discussion

The BCI system presented as part of this work represents one of the firsts of its kind to be designed for real-time speech-brain entrainment neurofeedback. Even though it was not possible to conduct the proposed experiment at the time this is being written, we have tested that our toolbox is capable of providing accurate and reliable real-time speech entrainment neurofeedback with simulated data. Therefore, it is ready to be used in actual experimental settings.

Our toolbox has minimal delays in all aspects. From the 2 ms delay in data collection into the buffer through *rda2ft*, to the average 218.9 ms in terms of PLV calculations. This maximizes the sense of agency for the participant and, consequently, BCI control as described by Evans et al., (2015). This is essential in order to develop strategies for top-down modulation of entrainment within auditory regions (Park et al., 2015). Moreover, our toolbox allows for the possibility of compiling it into a standalone application for users with a limited programming background and to further reduce the computation delays.

Nevertheless, the most remarkable feature of the toolbox is that it is built to be flexible and to allow the user to select various hyperparameters such the neurofeedback presentation windows or the frequency bands of interest depending on the experimental question. For example, in terms of the frequency bands of interest, the default theta/delta PLV average metric was picked to cover the spectrum of past studies highlighting the role of theta and delta brain oscillations in speech-brain entrainment (Bourguignon et al., 2013; Molinaro & Lizarazu, 2018; Lizarazu, Lallier & Molinaro, 2019). We believe that this default may be an appropriate metric to begin testing our toolbox in settings such as our proposed speech-in-noise validation experiment. Moreover, it could be used to test the effects of our neurofeedback technique on disorders like dyslexia given the atypical neural entrainment found in studies such as Lizarazu et al. (2015) and Molinaro et al. (2016). However, this default could be too broad and fail to capture specific entrainment features of either the delta or theta band for a given research question. In this vein, a researcher may choose to modify this default within the *RT_Entrainment_Toolbox_Main.m* script and elect to track just entrainment to the delta band by setting the frequency inputs to just the delta band instead of theta and delta for instance. In addition, for posterior clinical applications a user may even choose to track an entirely different frequency band such as the gamma band, which has been related to phoneme entrainment. This is relevant since phonemic oversampling has also been proposed as a potential cause of the phonological processing impairment characteristic of dyslexia (Giraud & Poeppel, 2012).

This level of flexibility within our novel neurofeedback technique thus opens up the possibility of several experimental paradigms that could become effective therapies for disorders such as the beforementioned dyslexia. Already, EEG-NF in general has already been shown to be successful in treating neuropsychological disorders ranging from ADHD (Sonuga-Barke et al., 2013) and epilepsy (Tan et al., 2009), to traumatic brain injury (Garzon, 2018) and autism spectrum disorder (Holtmann et al., 2011). Even in the case that speech-brain entrainment neurofeedback would be shown to be ineffective experimentally, the way we have built this toolbox also allows researchers and clinicians to swap the default audio comparison reference with an EEG channel of interest. Thus, making this toolbox relevant in all cases as it could apply traditional coherence-based neurofeedback, which has been already shown to be effective for epilepsy, autism spectrum disorder (ASD), and traumatic brain injury in previous studies (Coben, Wright, Decker & Morgan, 2015; Walker & Kozlowski, 2005; Rostami et al., 2017).

With that being said, this toolbox has its limitations. Some could be addressed with further development effort. Examples of these include the fact that it only supports BrainProducts technologies (BrainAmp amplifiers and BrainVision Recorder) along with the need for a very powerful hardware setup with a Linux operating system for the data processing and presentation computer. Moreover, the real-time data preprocessing currently implemented is limited. We rely on FIR band-pass filtering to attenuate high-frequency artifacts and trigger presentation pauses to control for actions such as blinking when there are no stimuli being presented. Real-time applications are harder to preprocess than an offline experiment without a major loss in latency. However, there are techniques that could be implemented like artifact subspace reconstruction (ASR) that would allow for thorough and efficient artifact rejection in real-time (Chang, Hsu, Pion-Tonachini & Jung, 2018).

Lastly, there is an inherent limitation with all neurofeedback applications, which is their underlying controversy regarding their reliability as clinical tools. There are study reviews that claim that the findings showing that neurofeedback applications are effective have several limitations that range from small sample sizes to the lack of study randomization (Micoulaud-Franchi, 2015; Schoenberg & David, 2014). However, the biggest limitation is the unclear long-term effects of neurofeedback therapies. In this regard, previous studies have found mixed results. For instance, Janssen et al. (2020) did not find any specific long-term benefits of theta/beta neurofeedback in the treatment of ADHD compared to pharmacological and physical treatment. On the other hand, Kouijzer et al. (2009) found significant long-term effects of theta/beta modulation neurofeedback in ASD patients. This level of variability makes it hard to predict whether speech-brain entrainment neurofeedback will have robust long-term effects. However, given how atypical entrainment appears to be present in disorders such as dyslexia as shown by previous studies (Hämäläinen, Salminen, & Leppänen, 2013; Lizarazu et al., 2015; Molinaro et al., 2016), it is likely that speech-brain entrainment can lead to reliable behavioral improvements. In any case, the long-term effects of speech entrainment neurofeedback along with the other limitations are certainly aspects that will be carefully looked at, and controlled for, in future studies looking to further evaluate our toolbox for clinical applications.

## Conclusion

Overall, despite the testing limitations, we have implemented an open-source BCI toolbox that is a robust piece of software capable of delivering real-time speech-brain entrainment neurofeedback with minimal delays in all fronts. There are well-founded reasons to believe that speech-brain entrainment neurofeedback will be effective given the relevance of neural entrainment to speech in disorders such as developmental dyslexia, stroke, and Broca’s aphasia. Thus, there is a great potential for this toolbox to become the core of a cutting-edge clinical intervention.

## Appendix A: Matlab Scripts for the Real-Time Entrainment Toolbox

### RT_Entrainment_Toolbox_Main.m

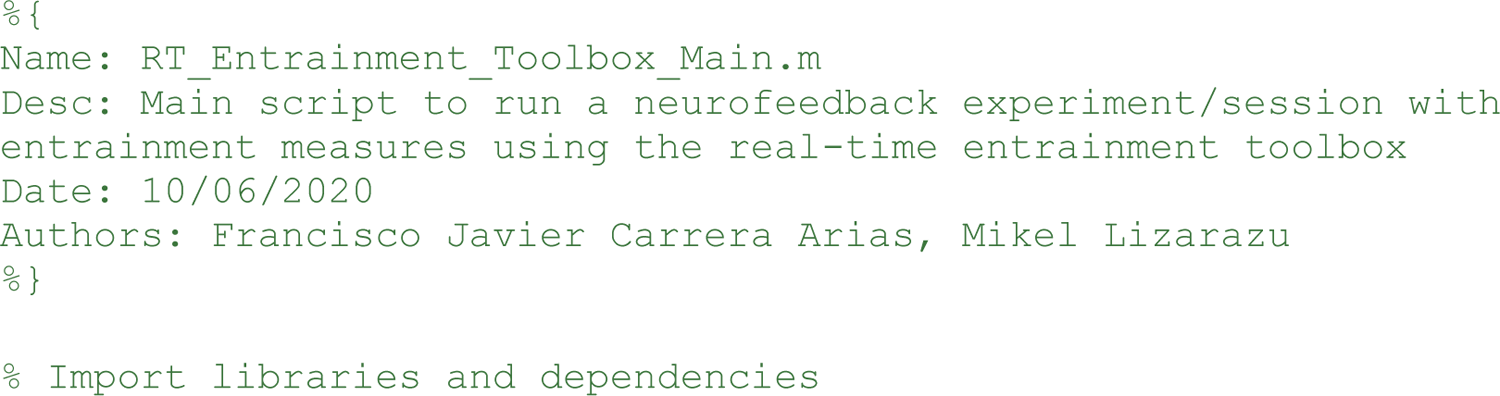
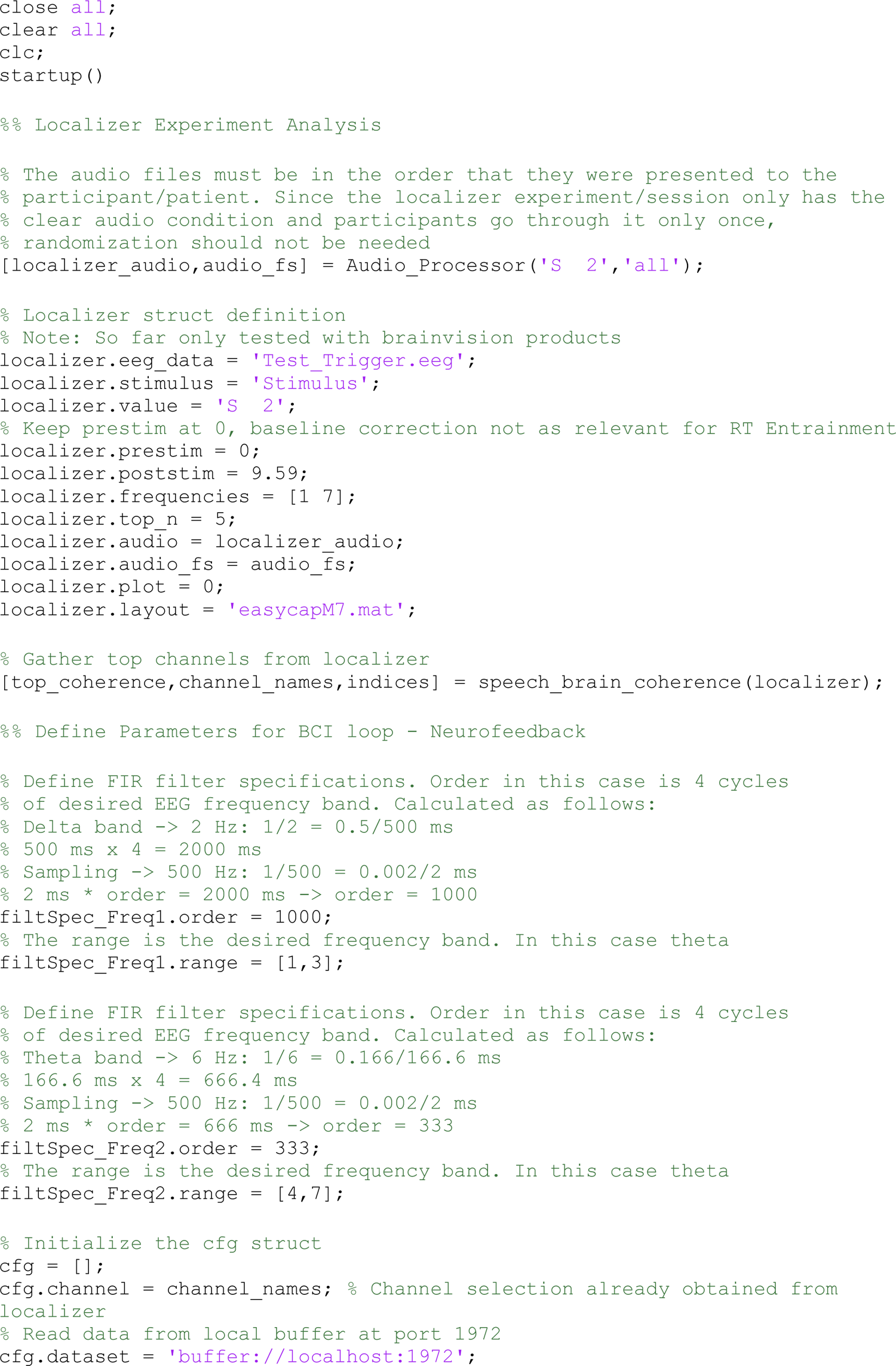
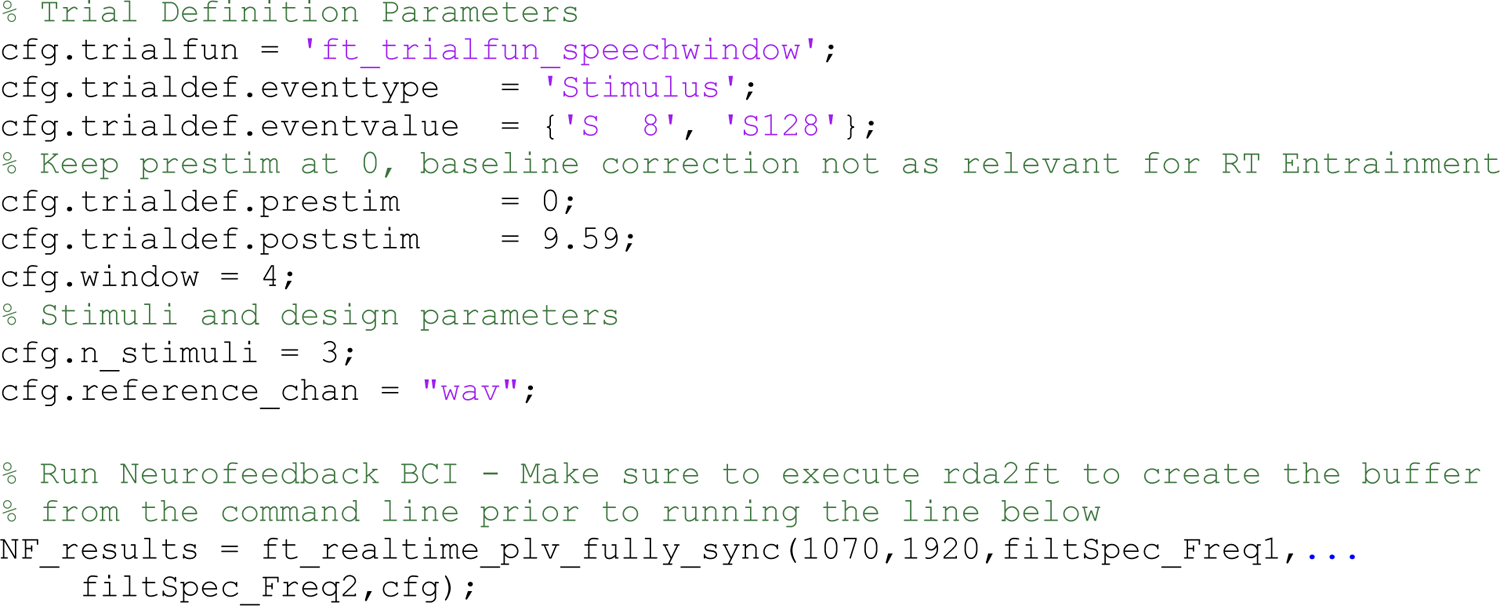

### Startup.m

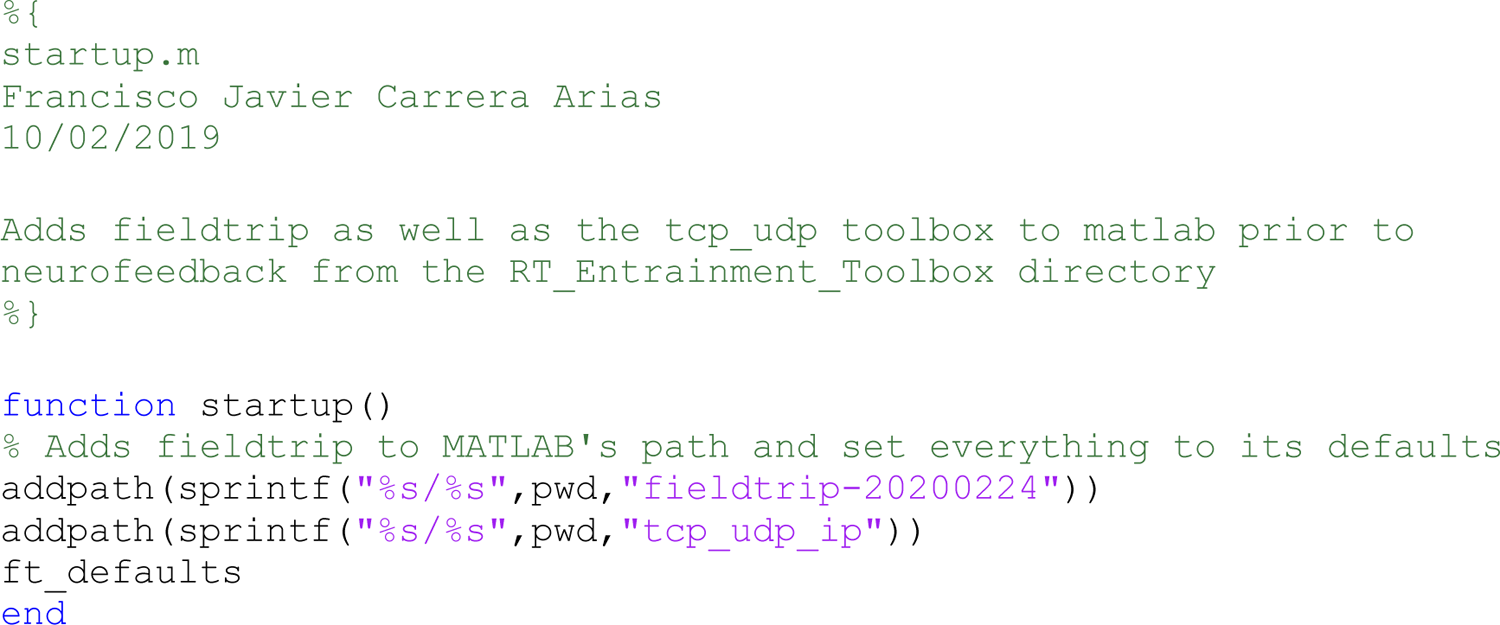

### Audio_Processor.m

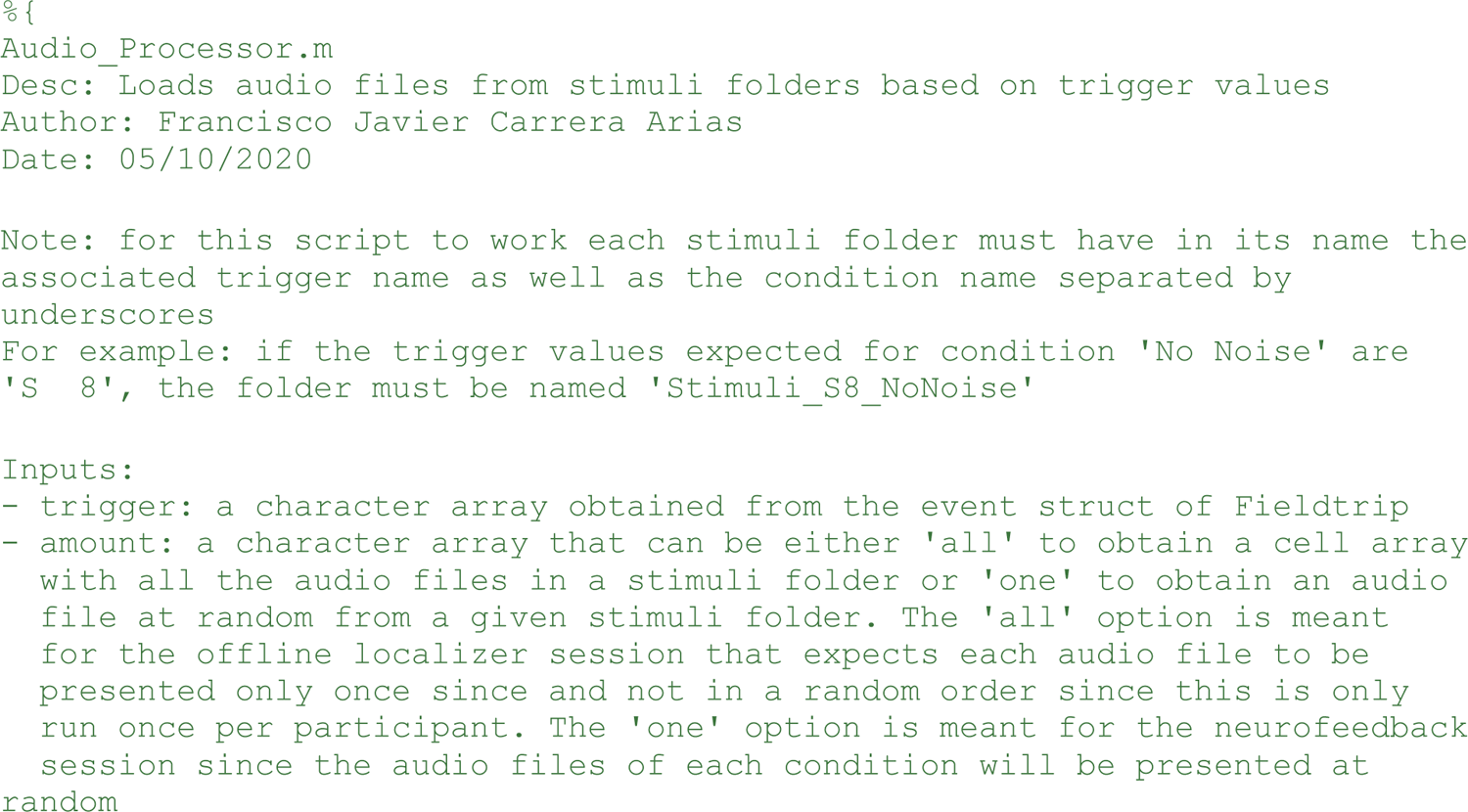
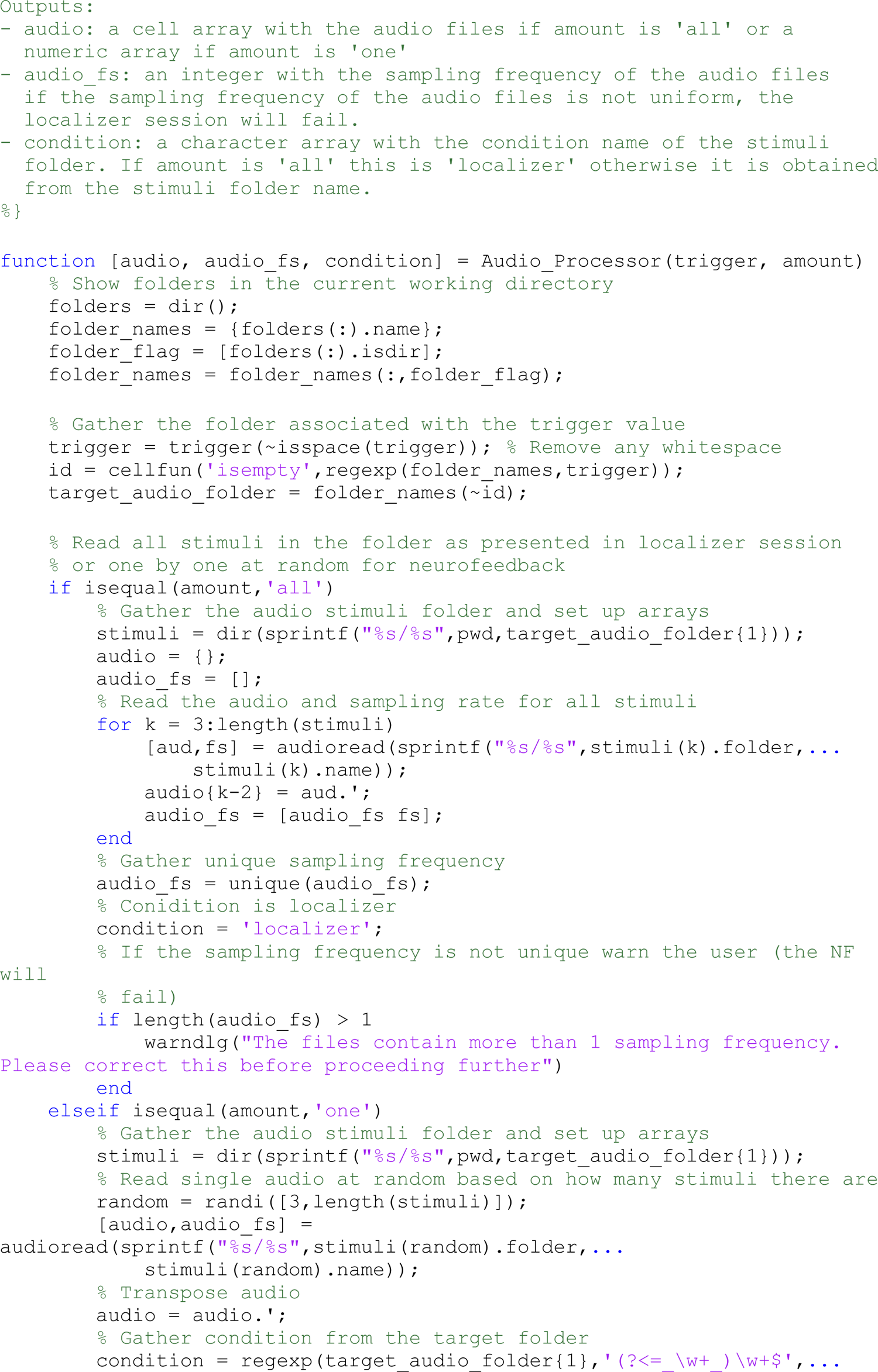
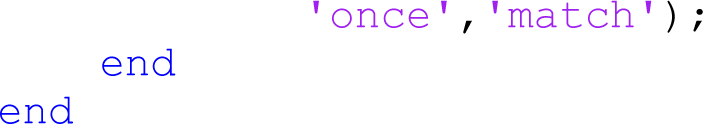

### Trial_Segment.m

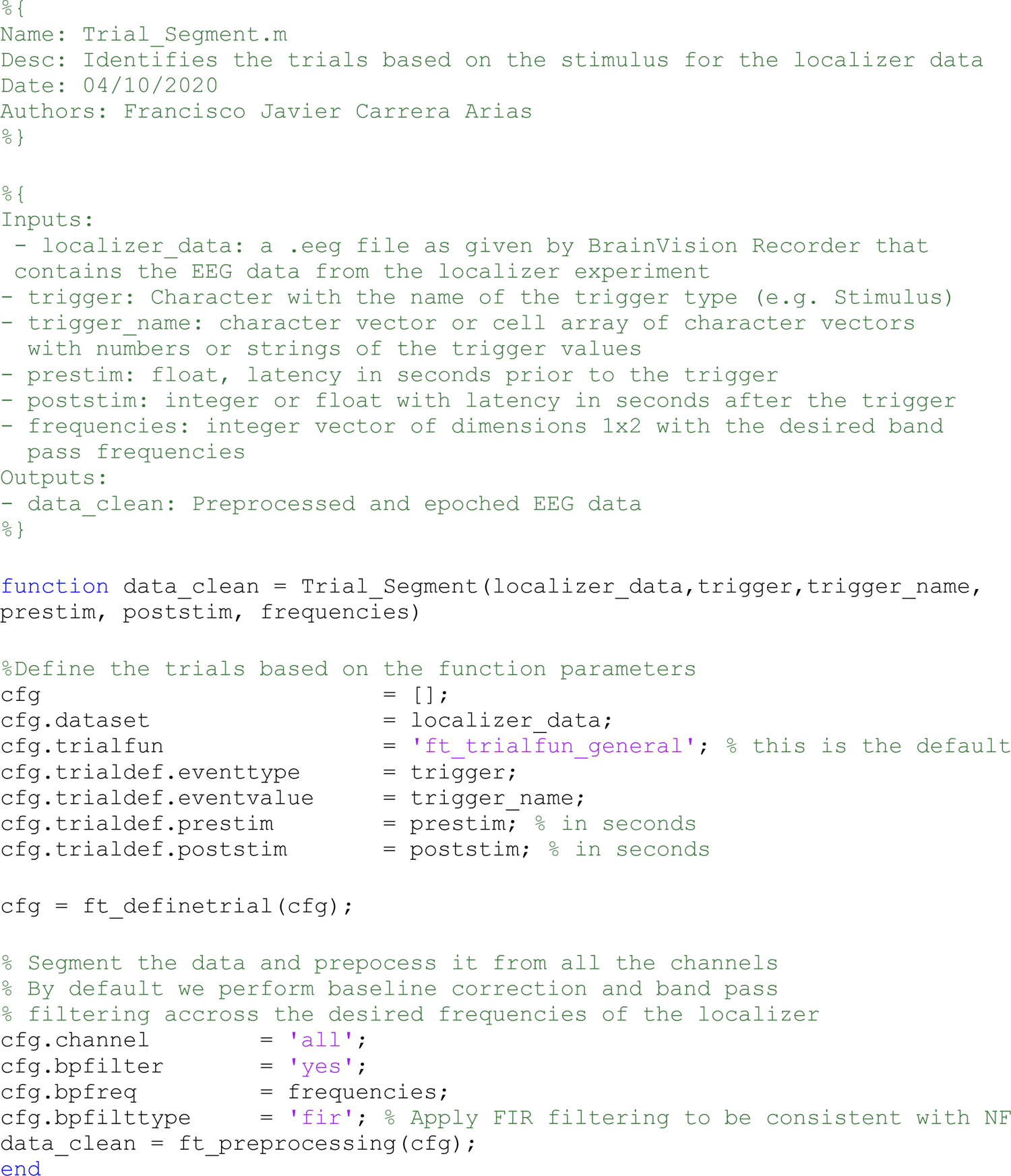

### audio_envelope.m

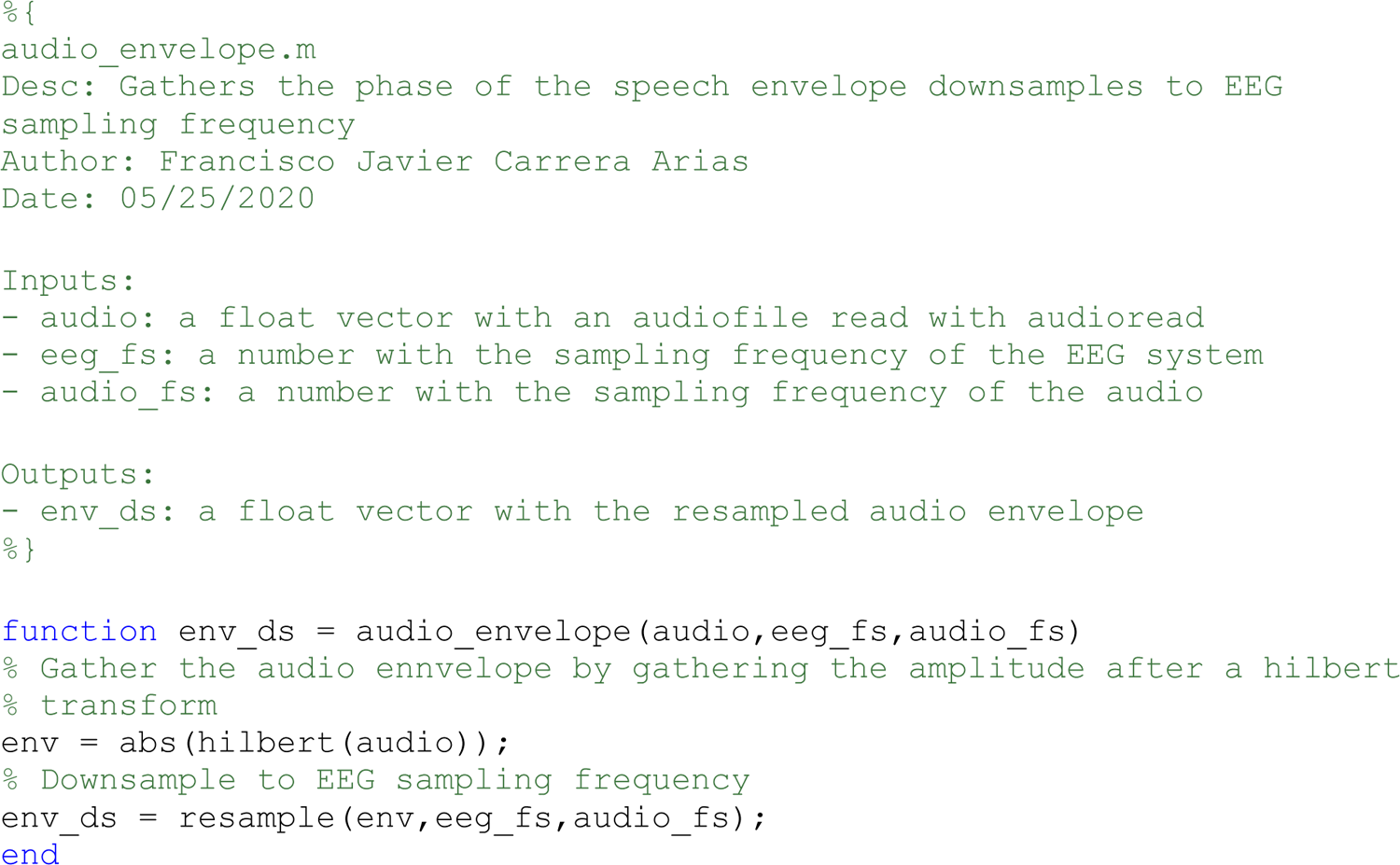

### speech_brain_coherence.m

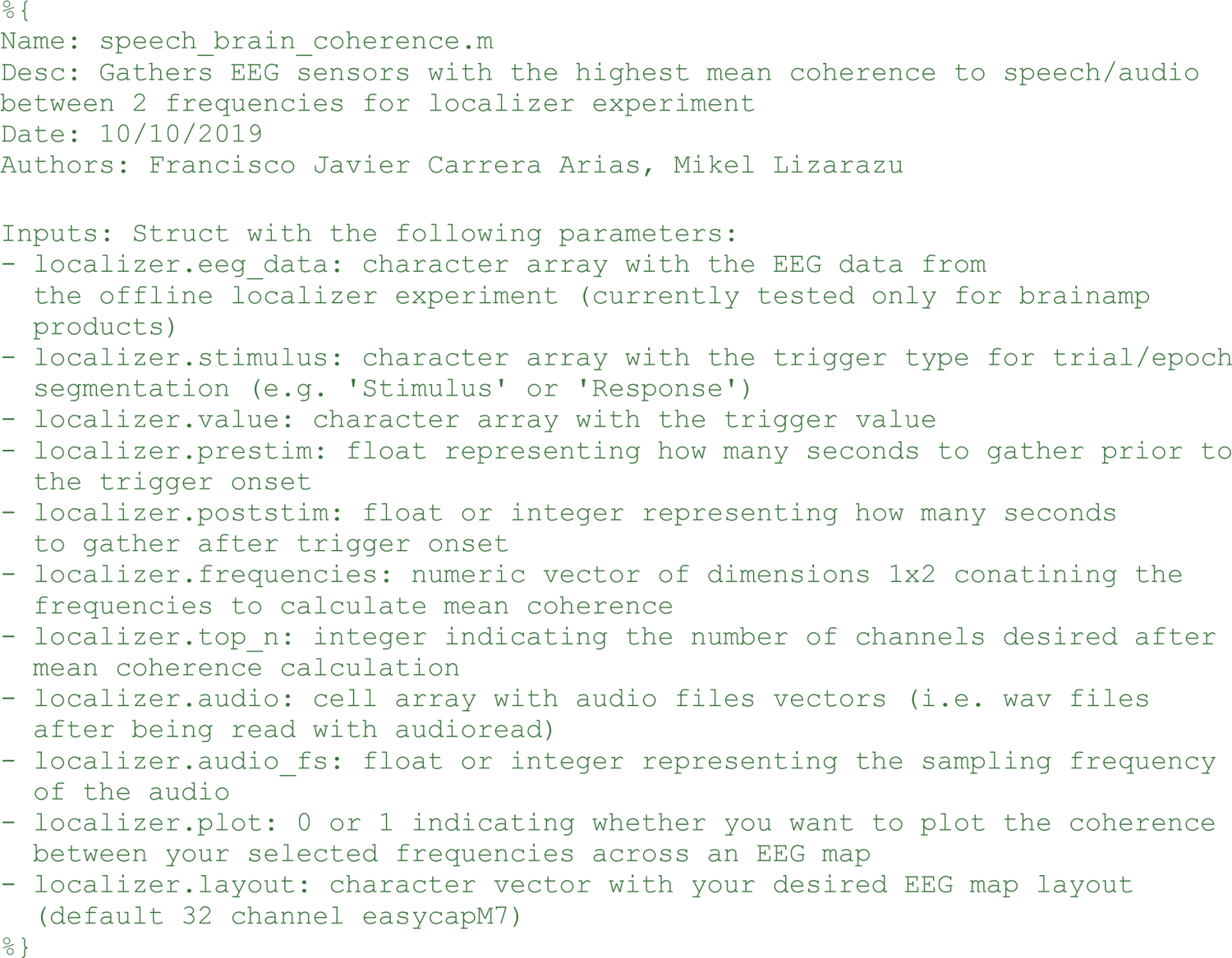
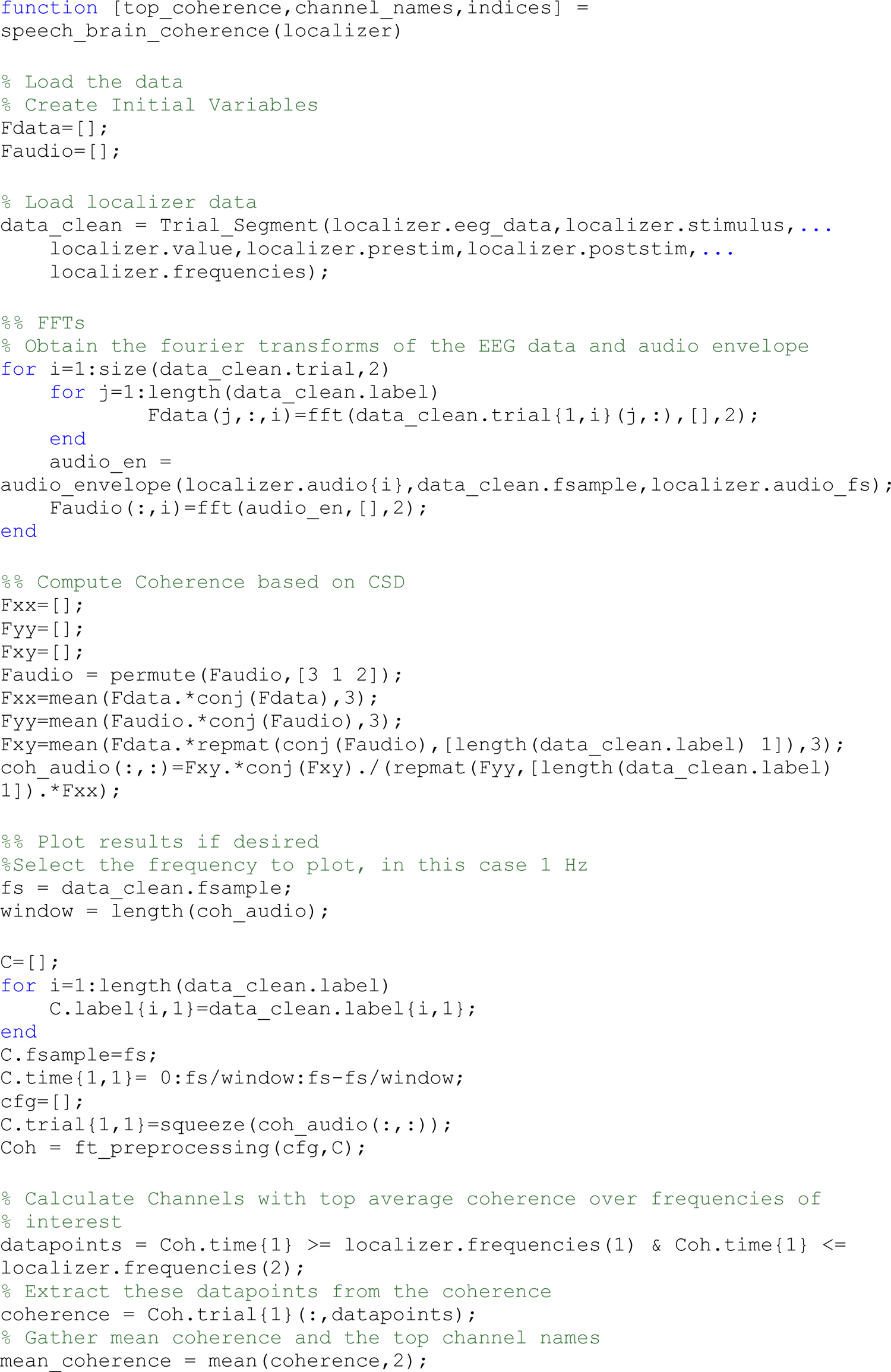
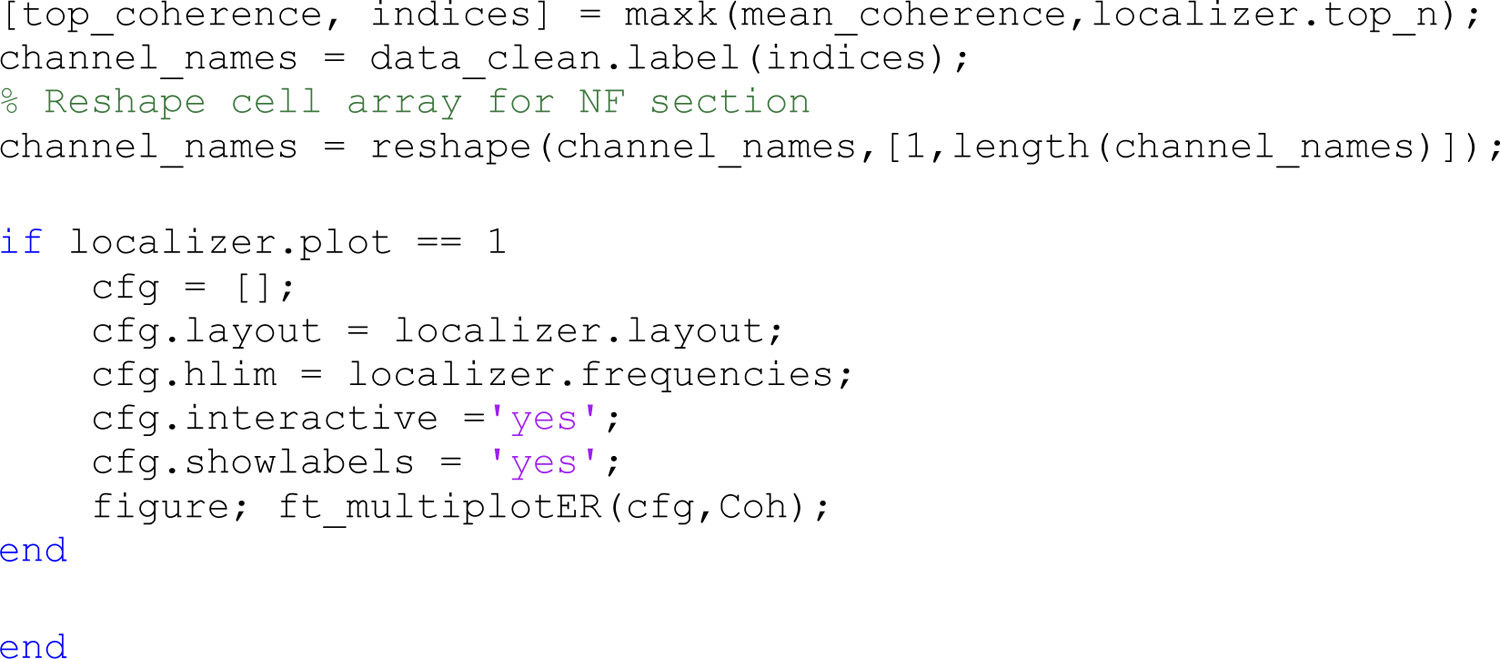

### Screen_Setup.m

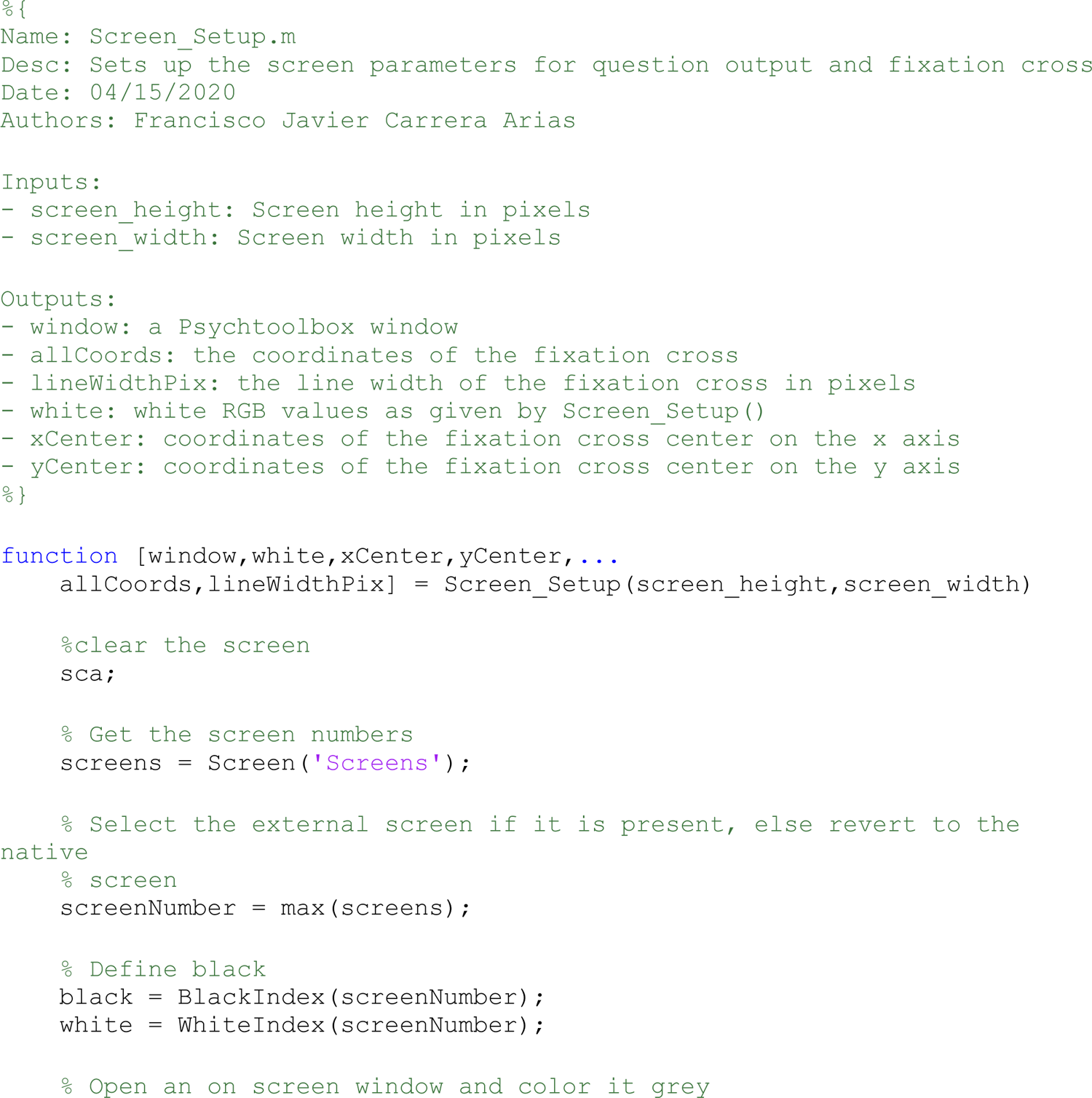
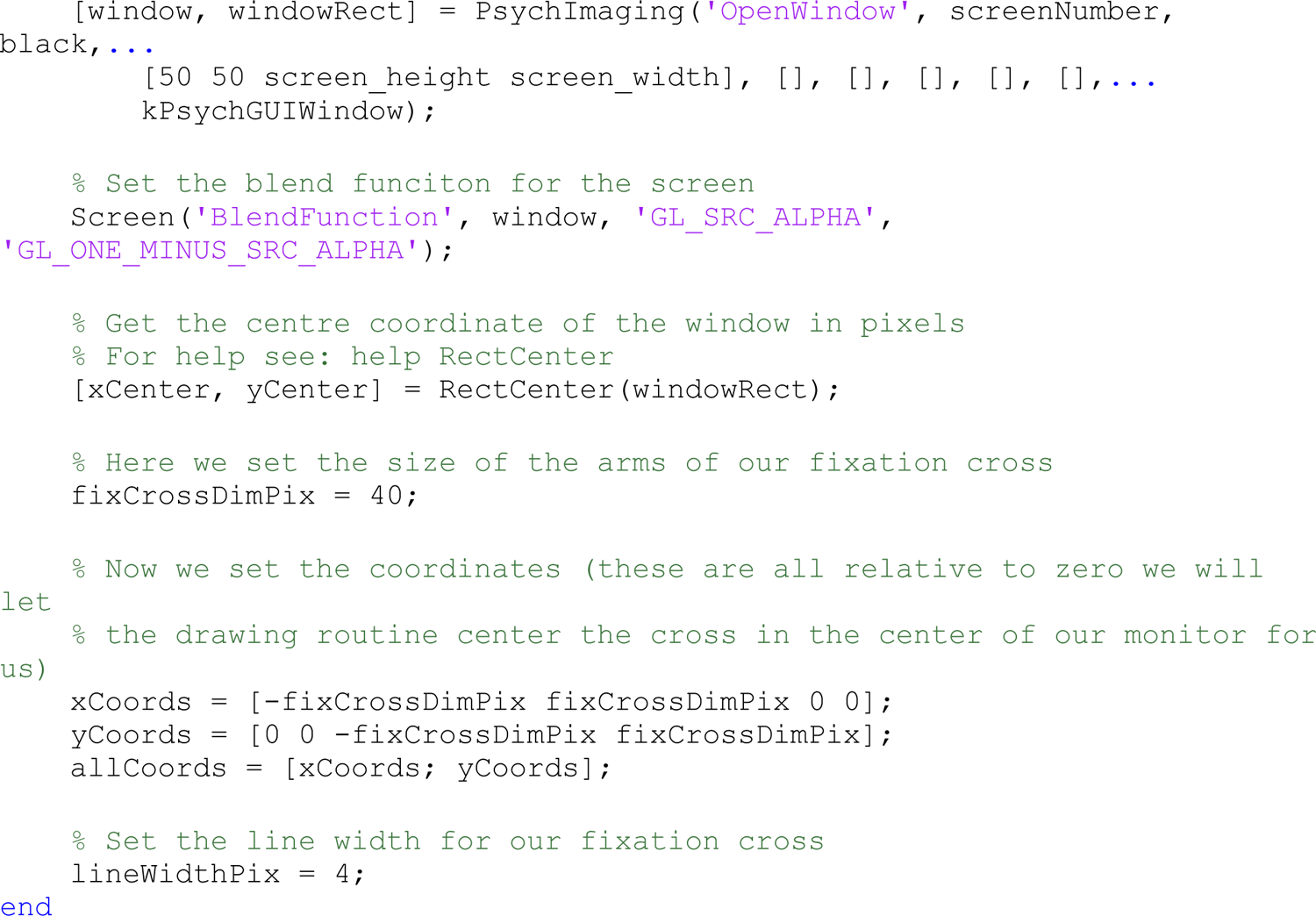

### Sound_Setup.m

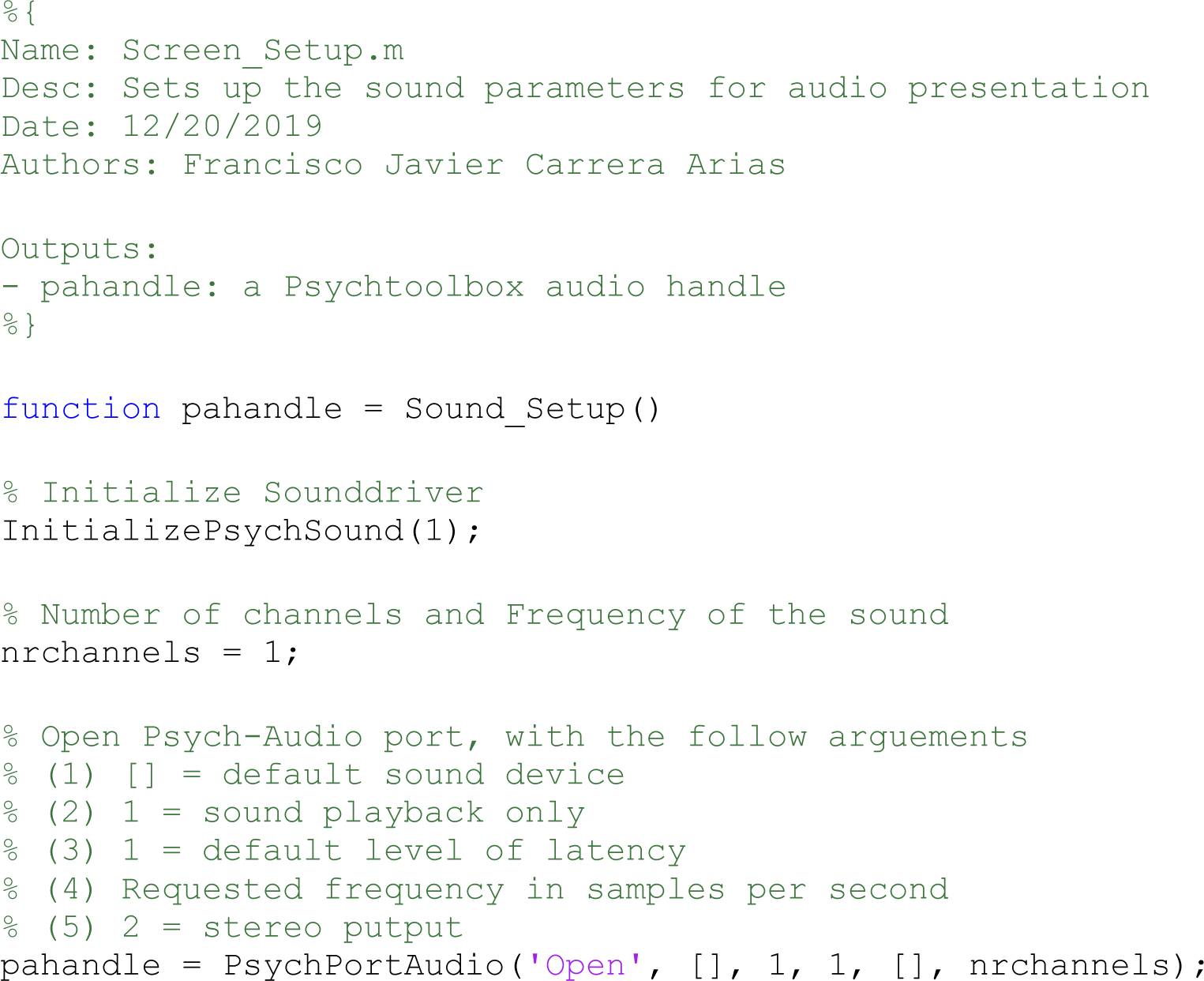
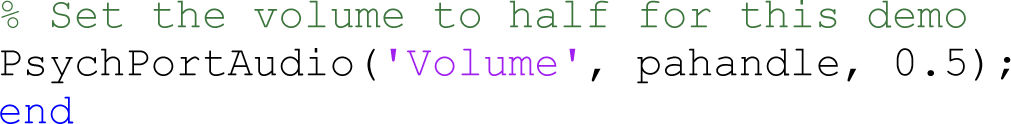

### Draw_Fixation.m

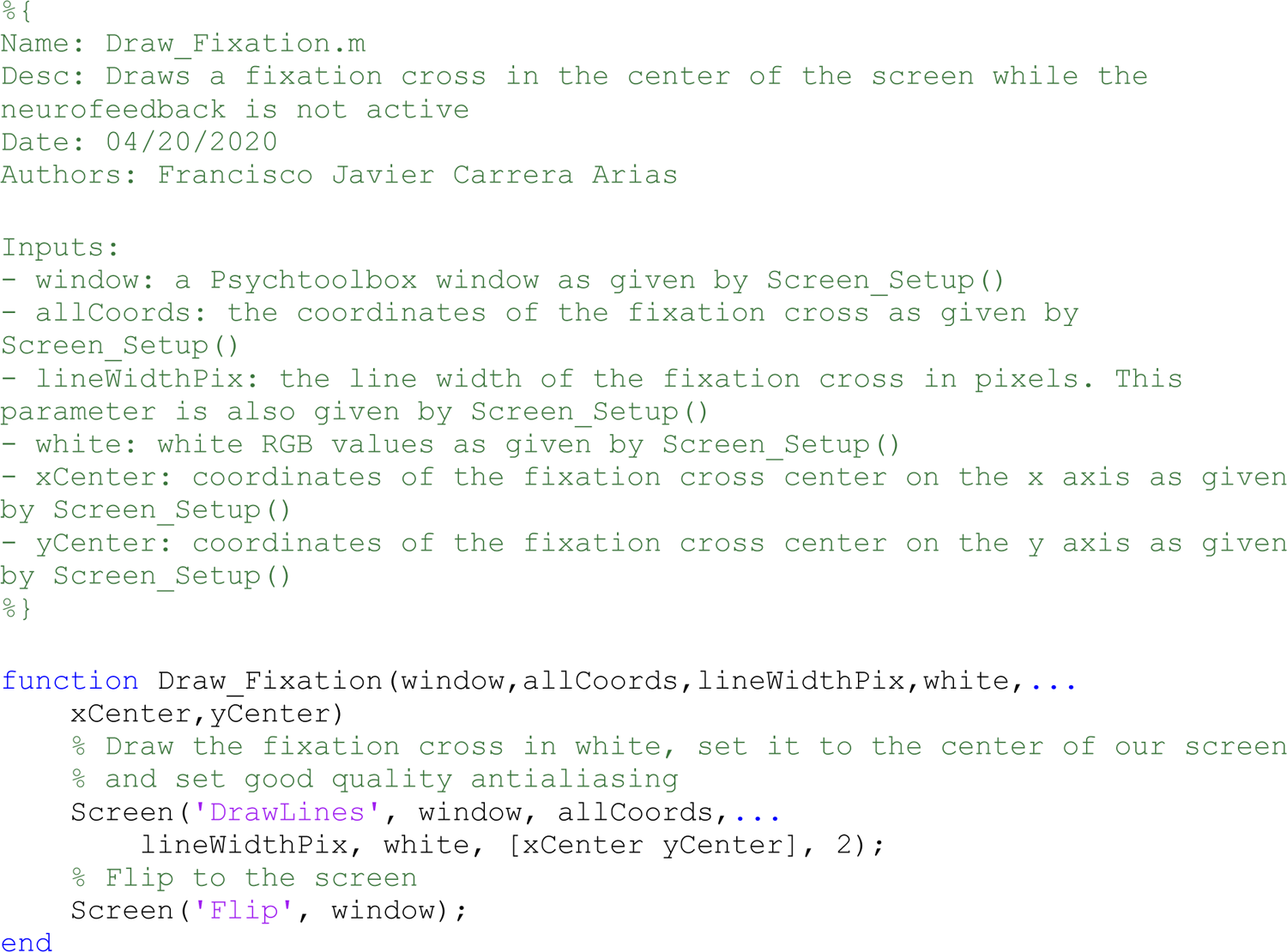

### ft_trialfun_speechwindow.m

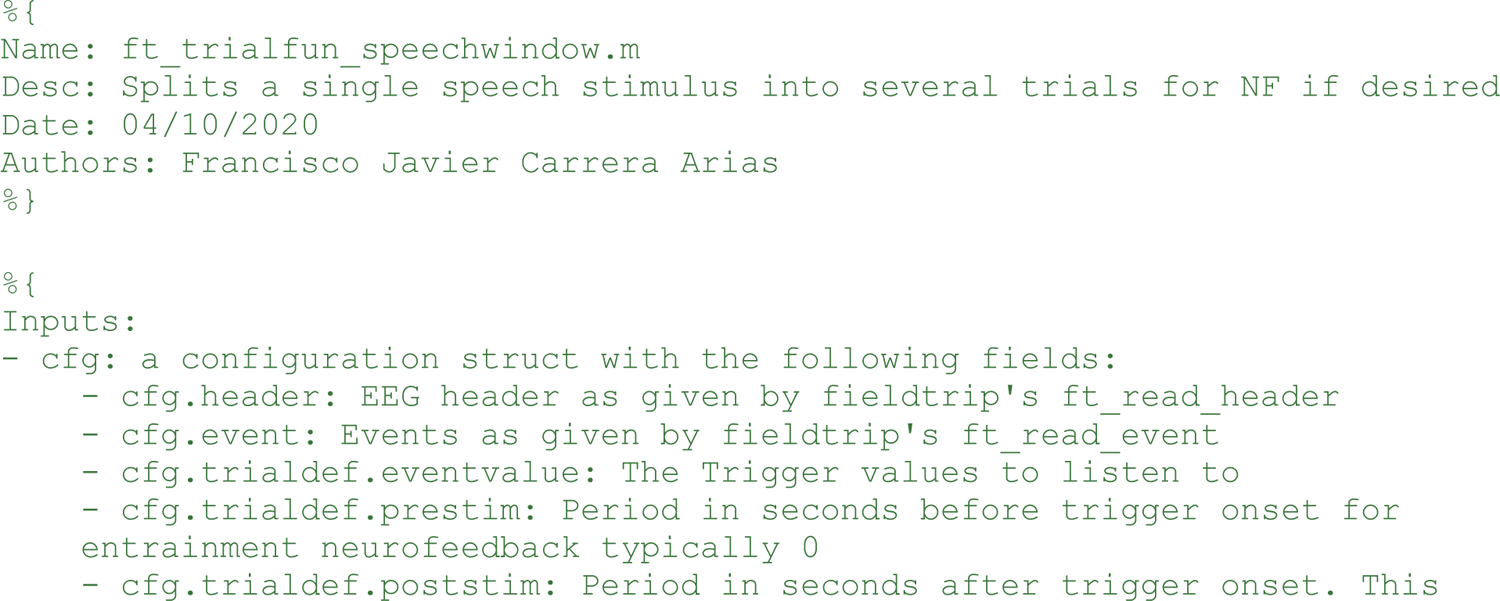
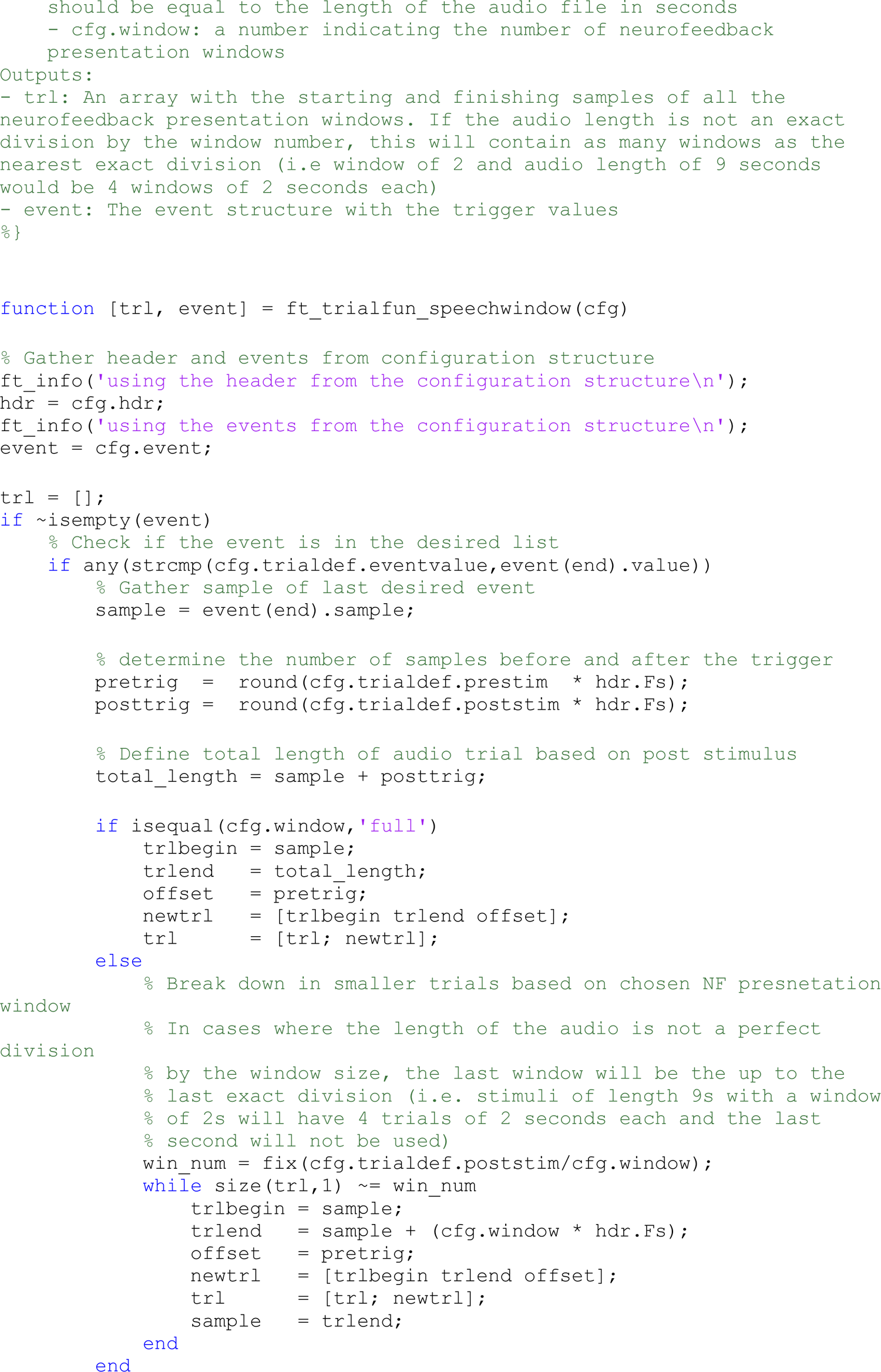
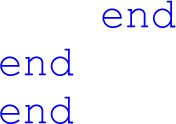

### plv_realtime_fun.m

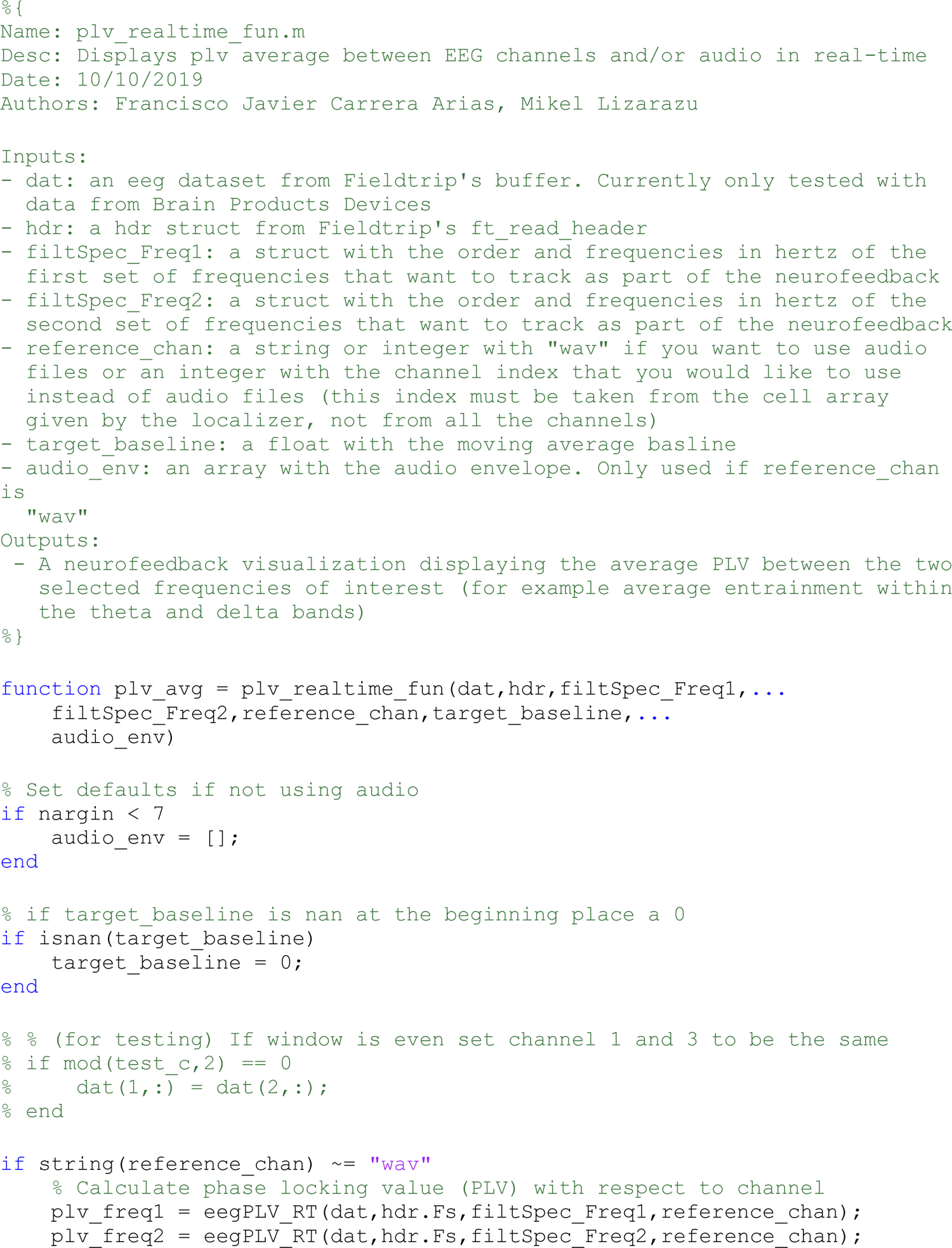
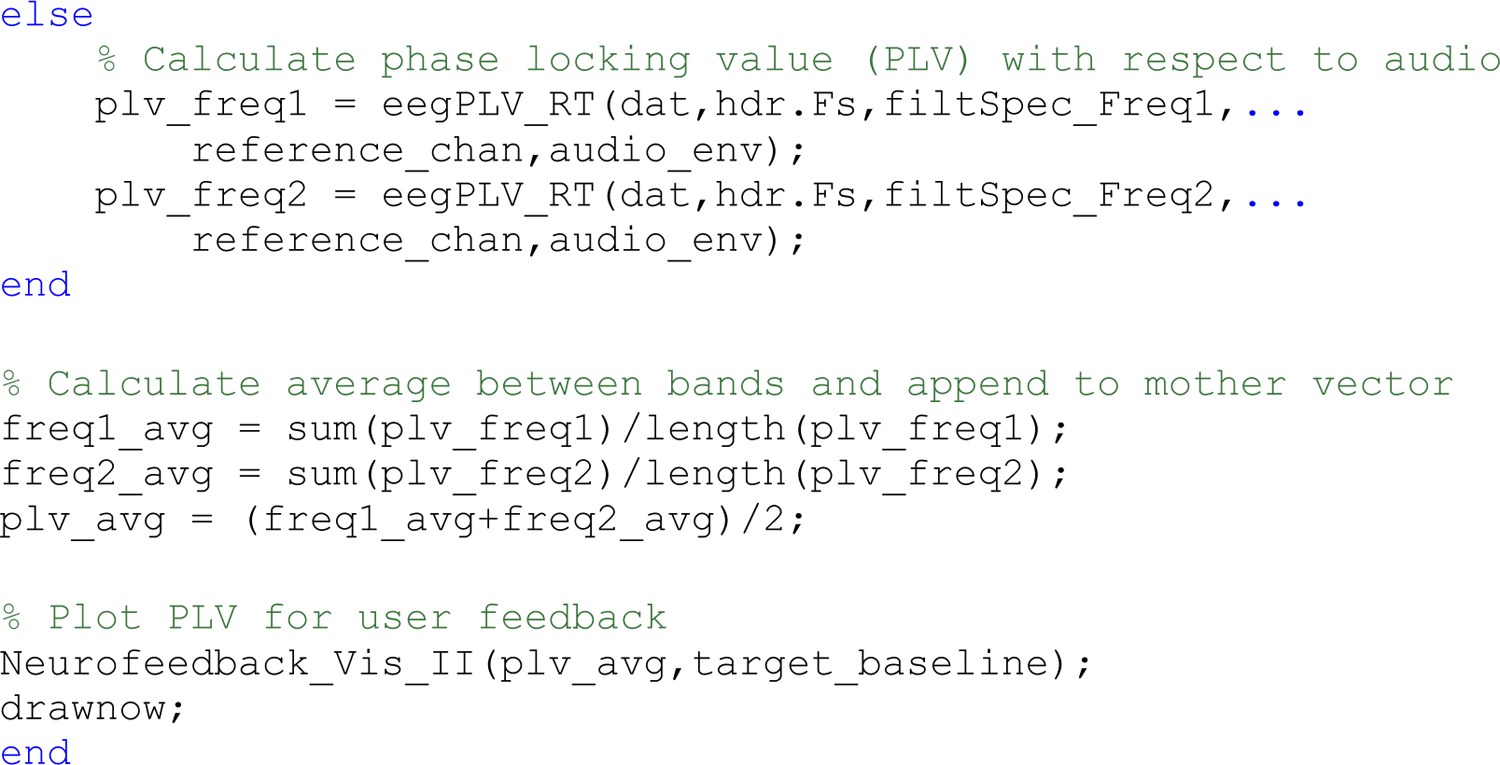

### eegPLV_RT.m

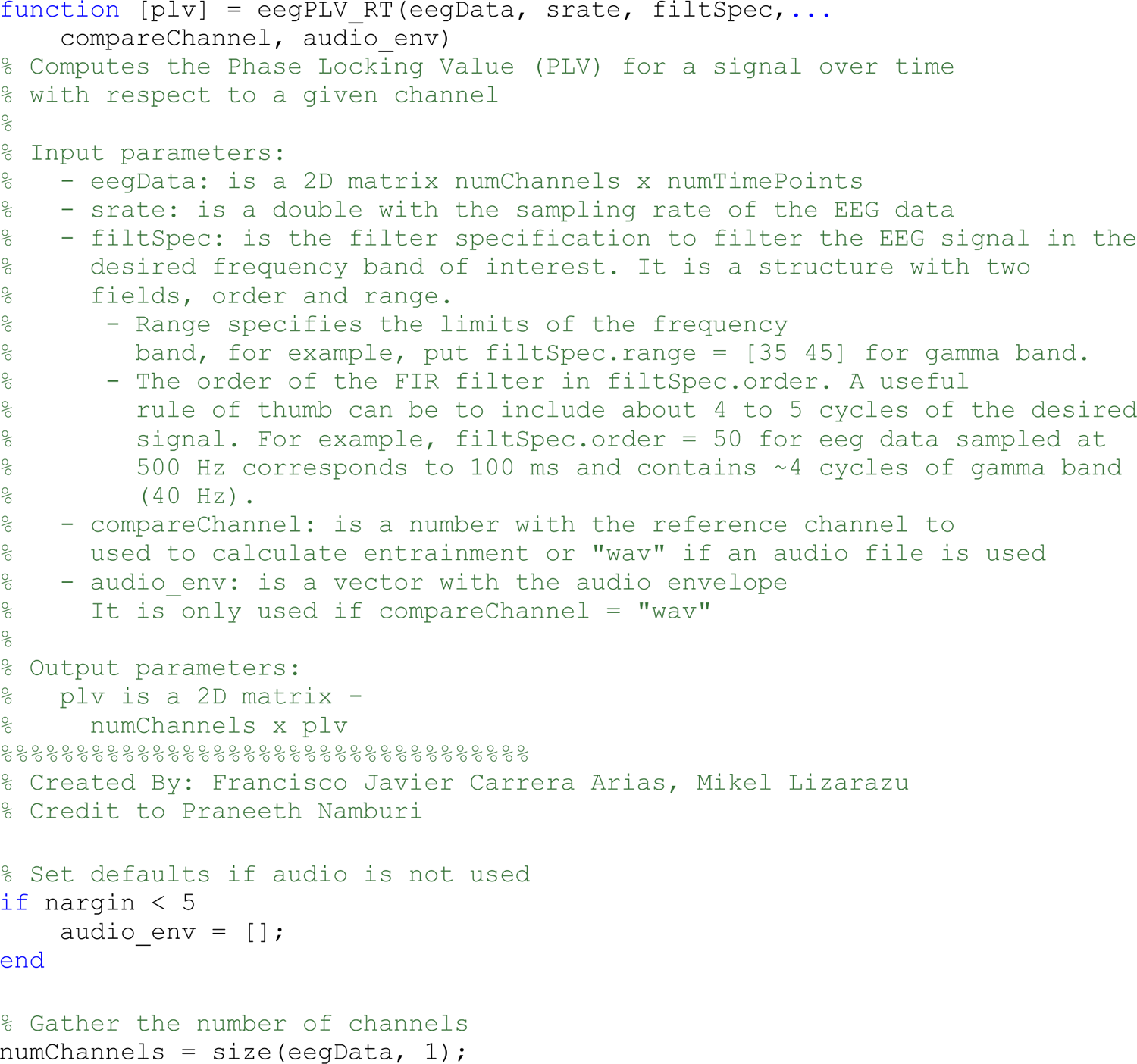
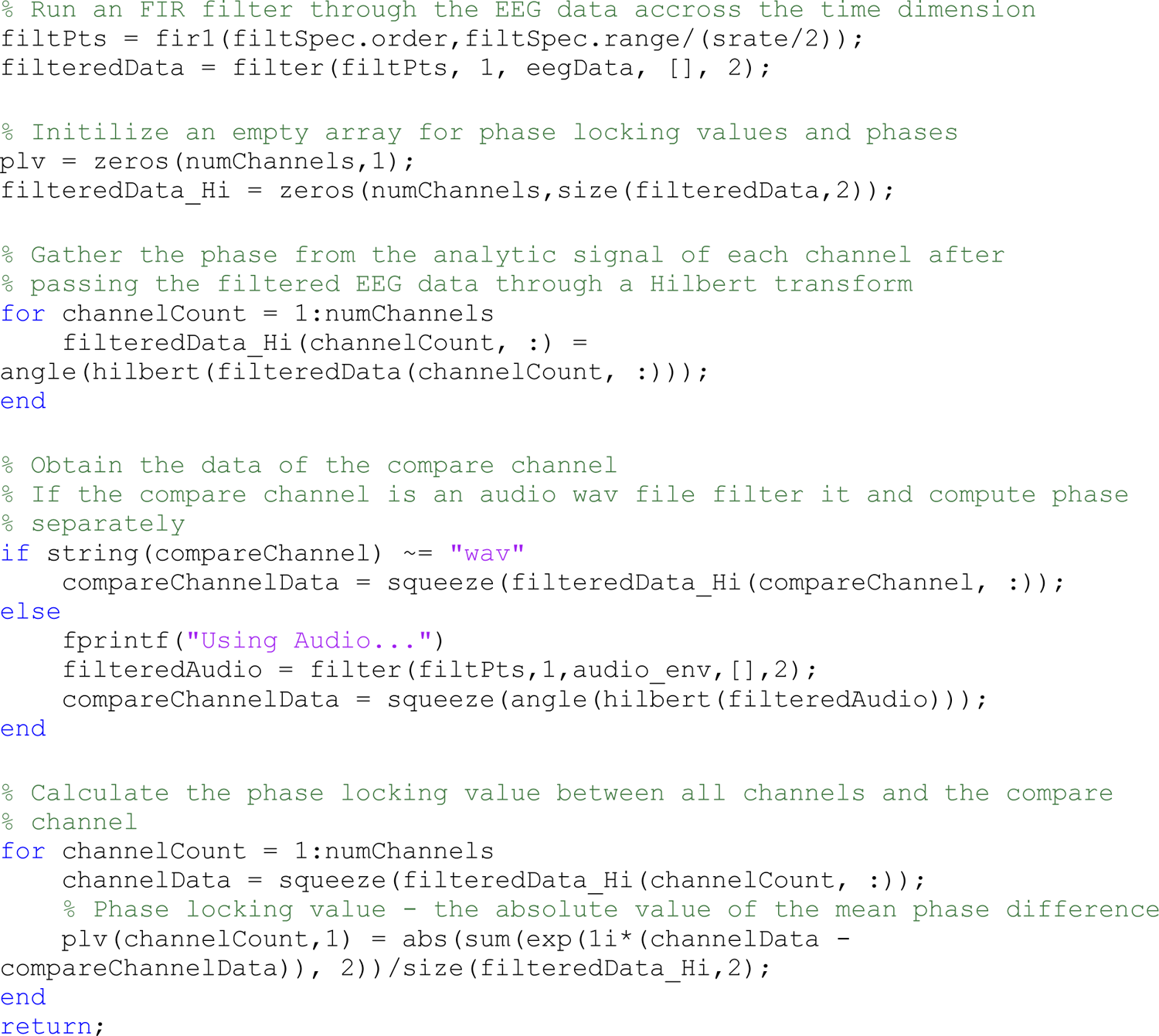

### Neurofeedback_Vis_II.m

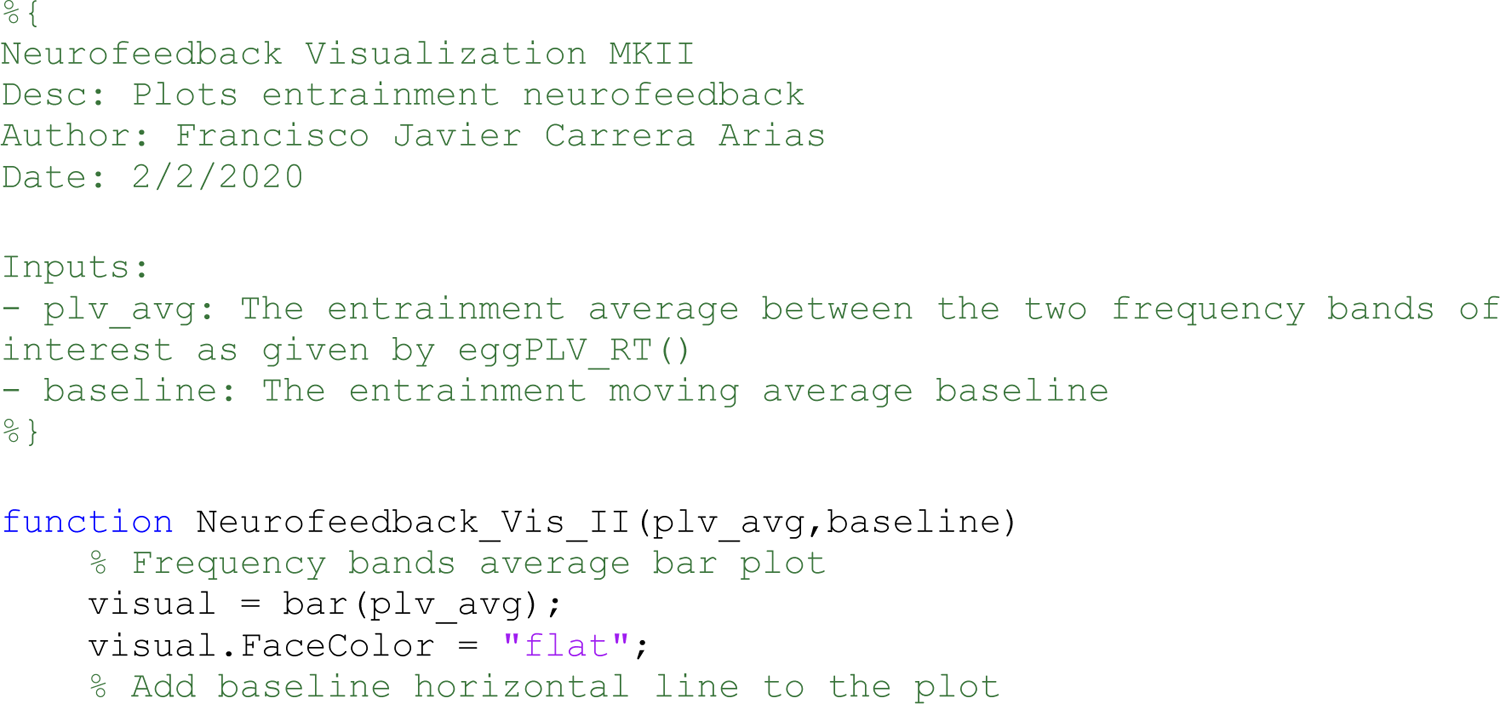
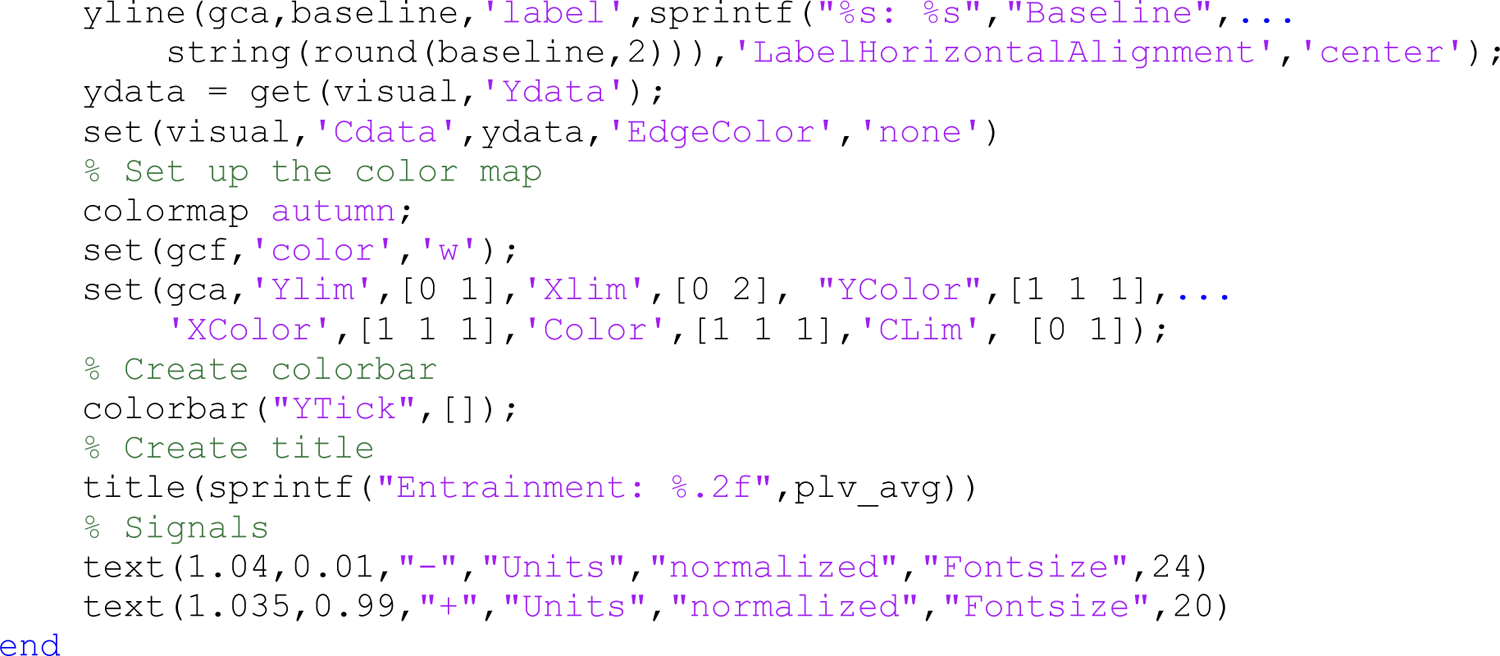

### Intel_Question.m

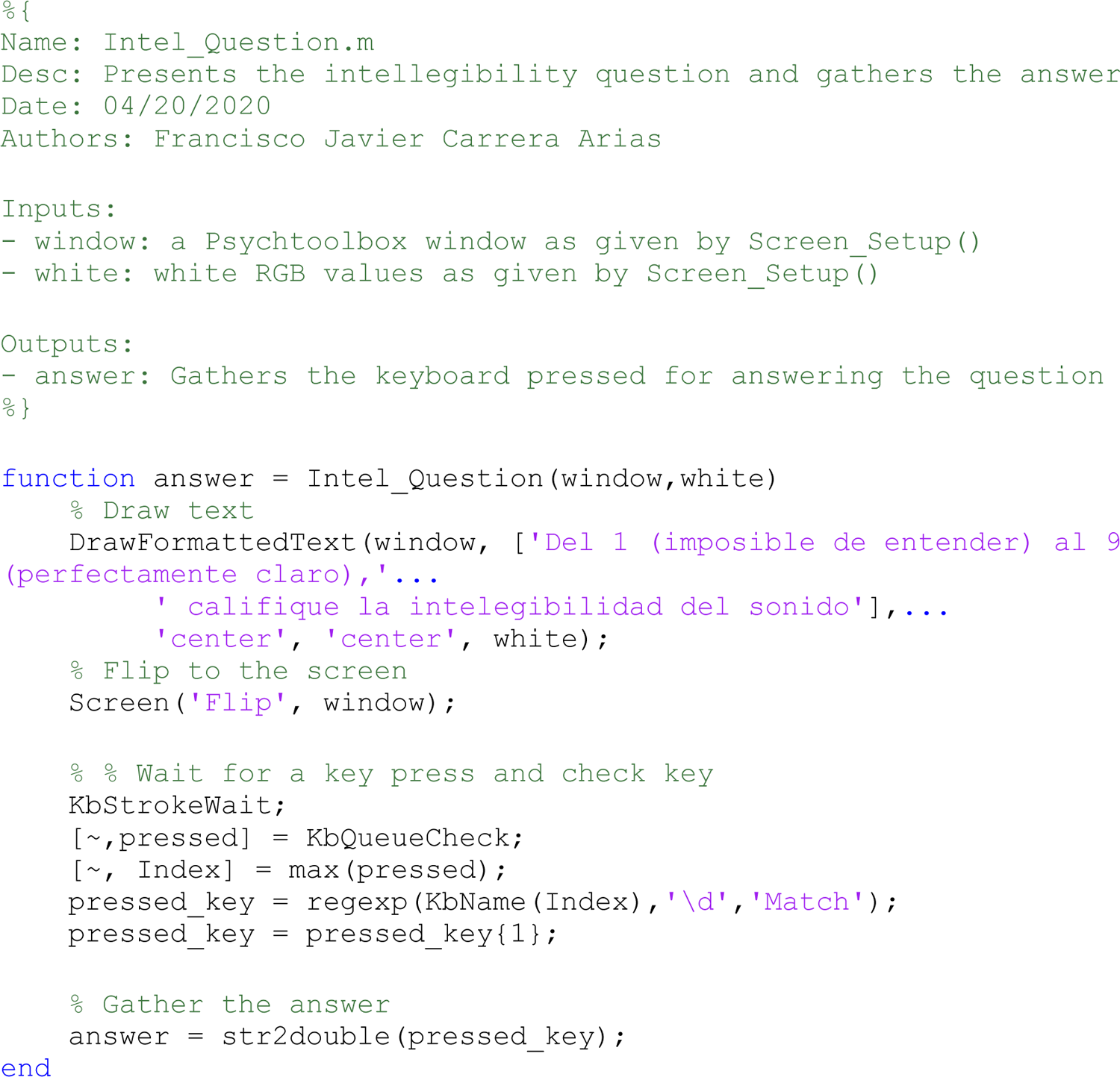

### ft_realtime_plv_fully_sync.m

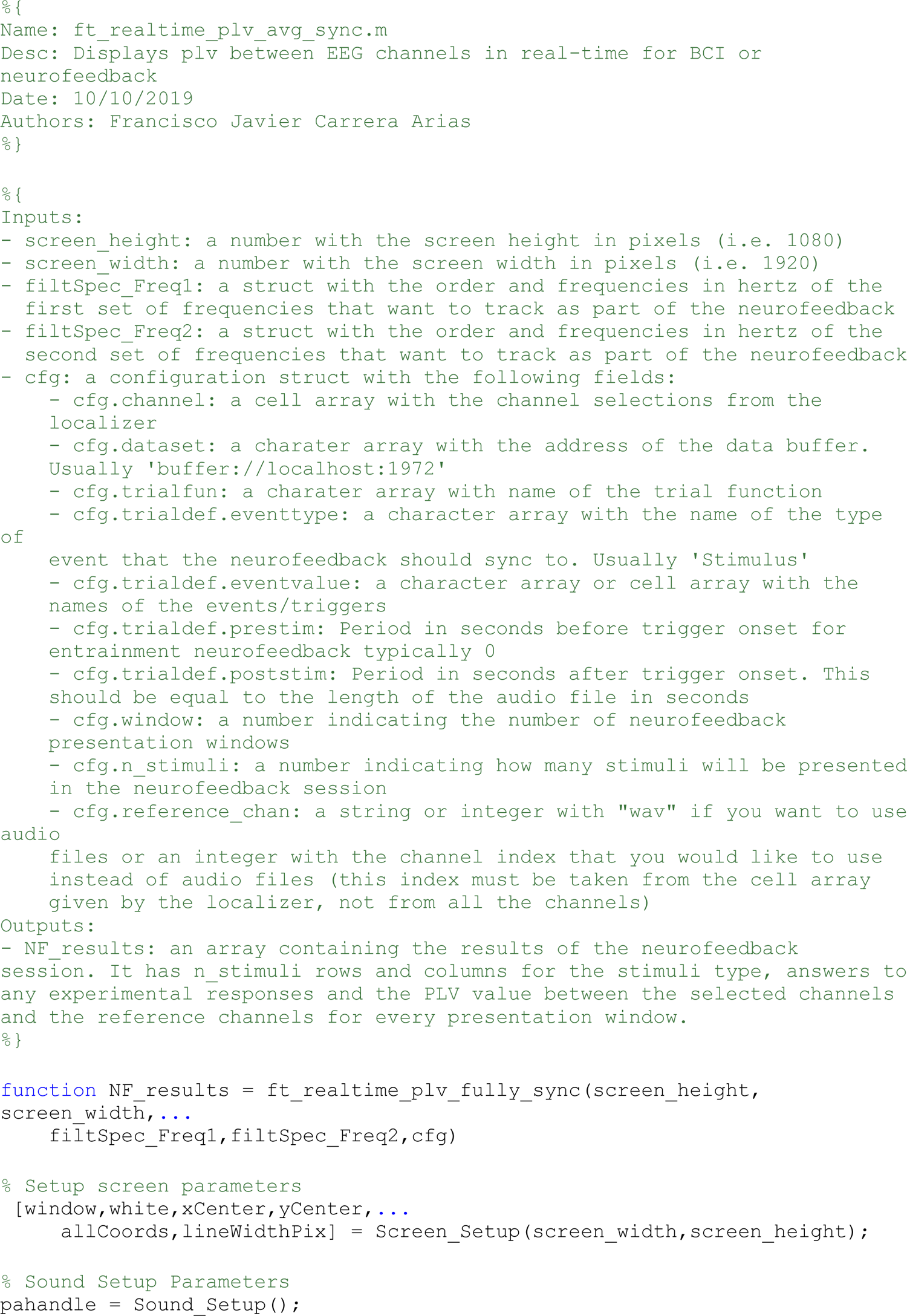
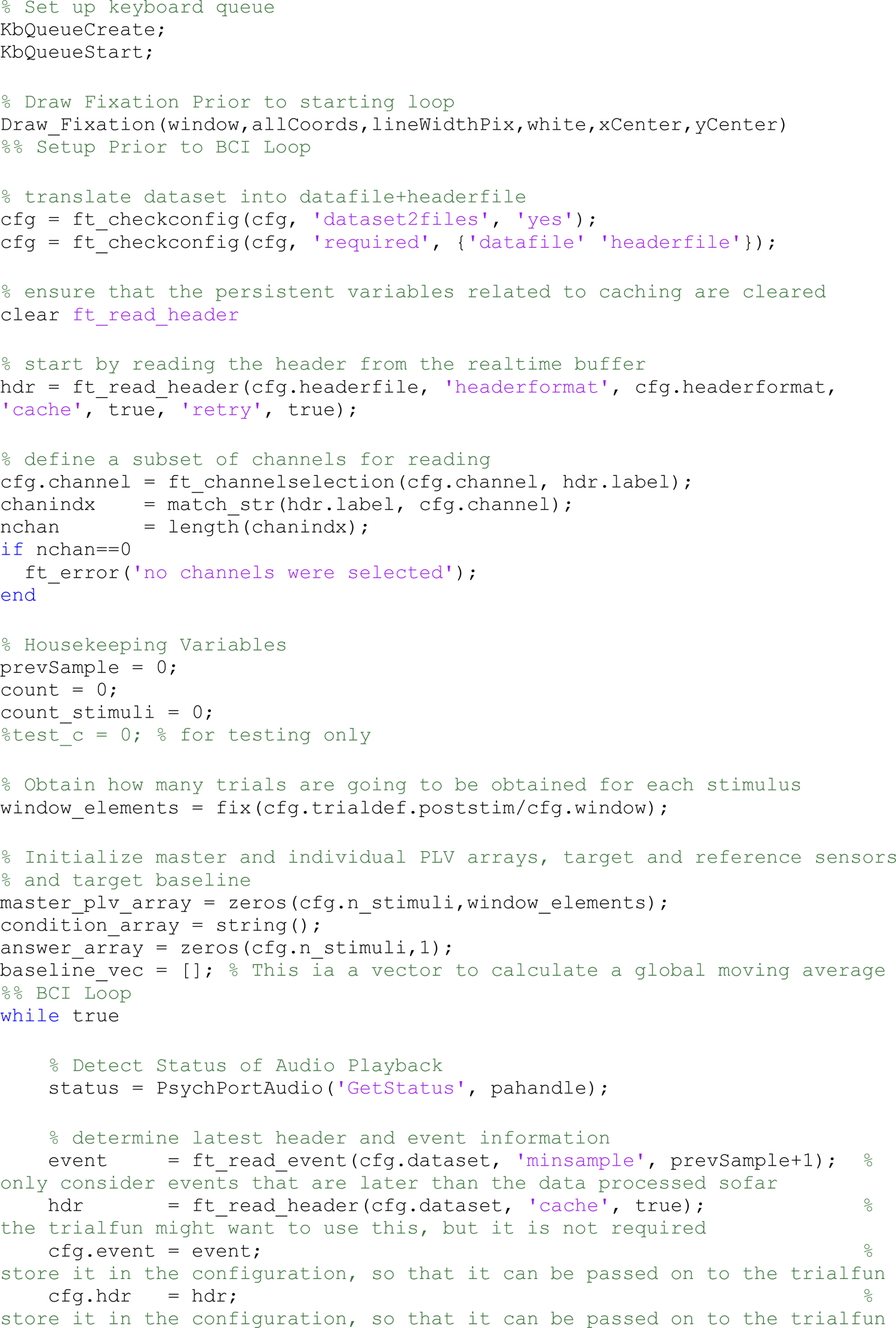
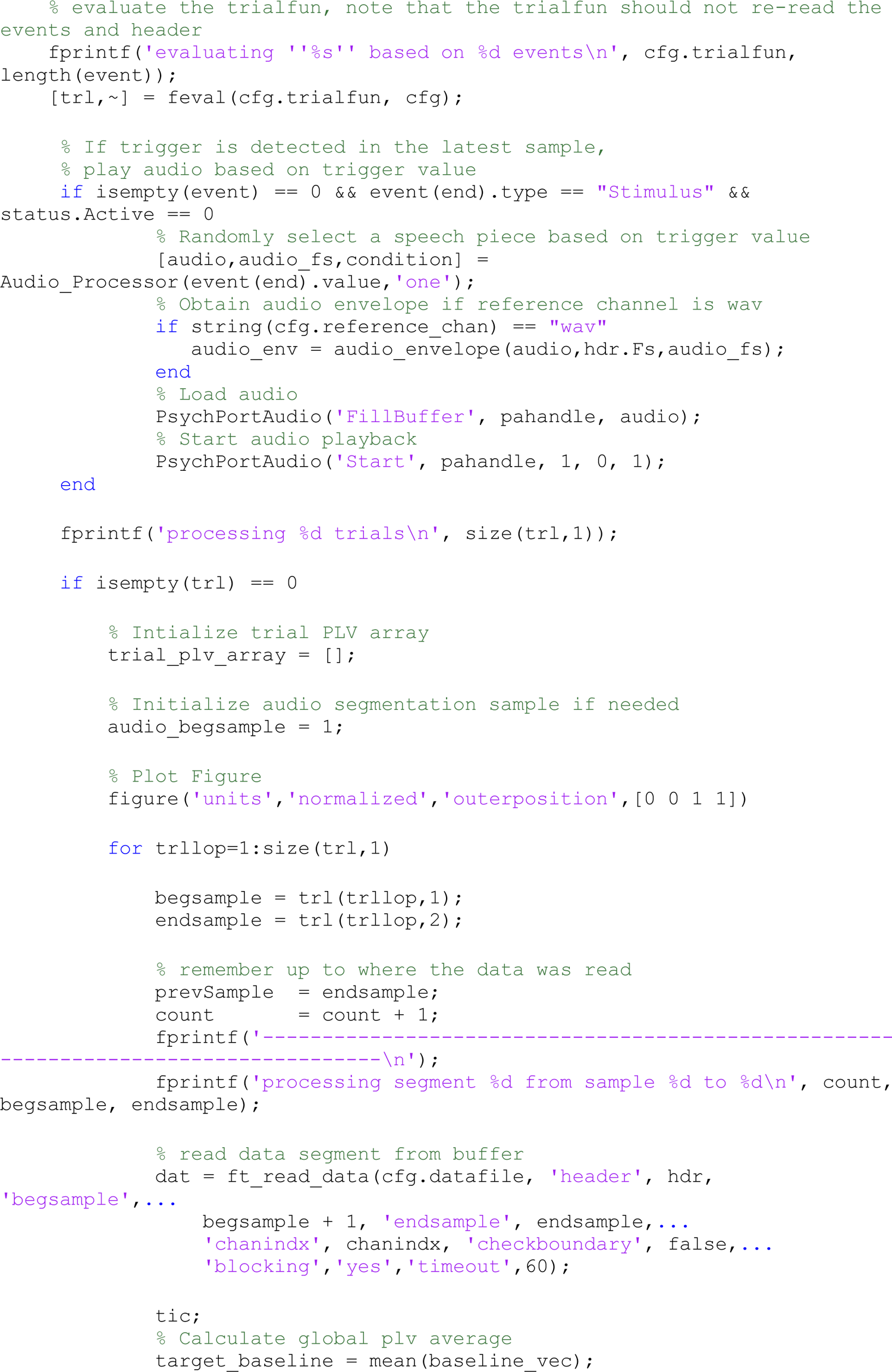
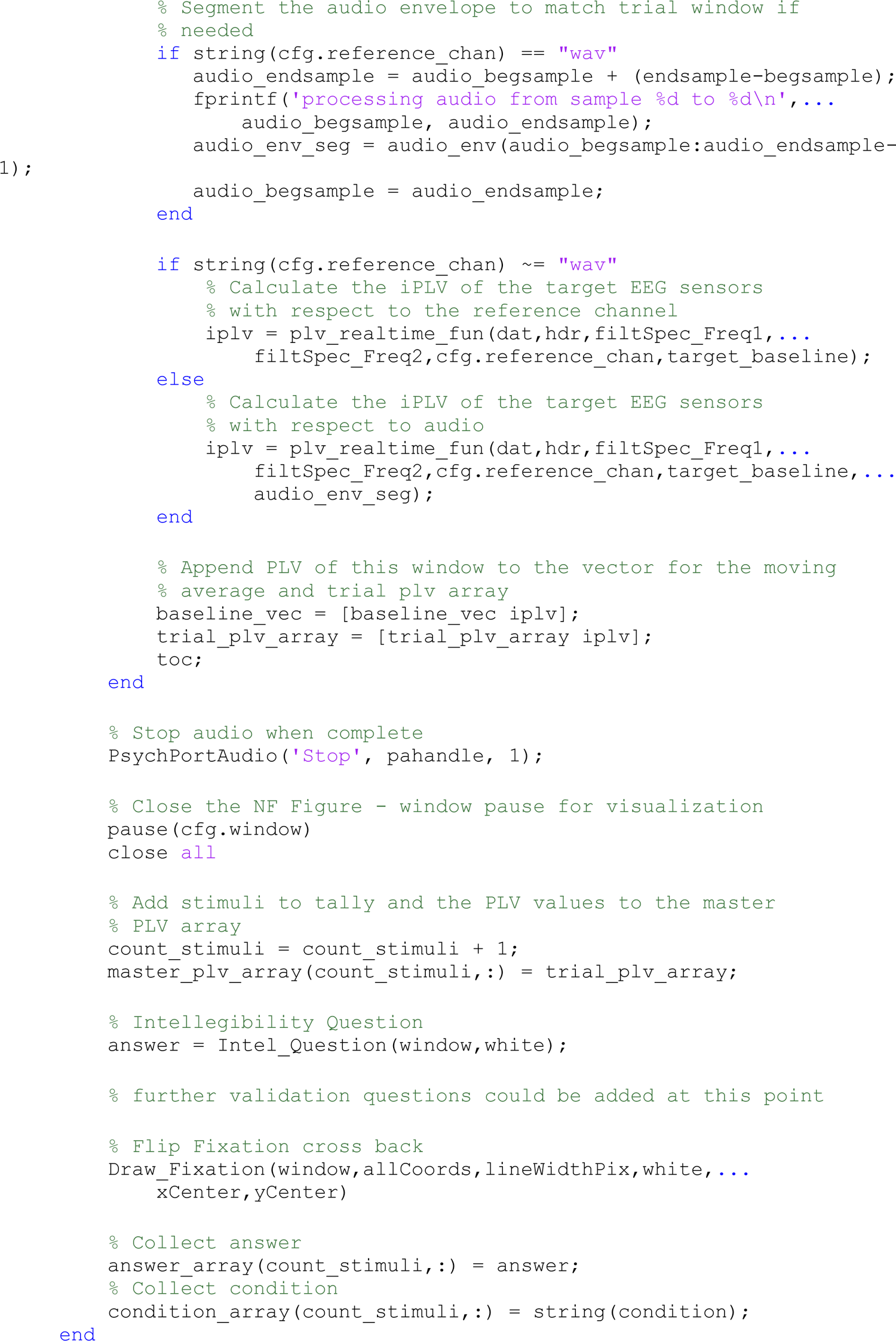
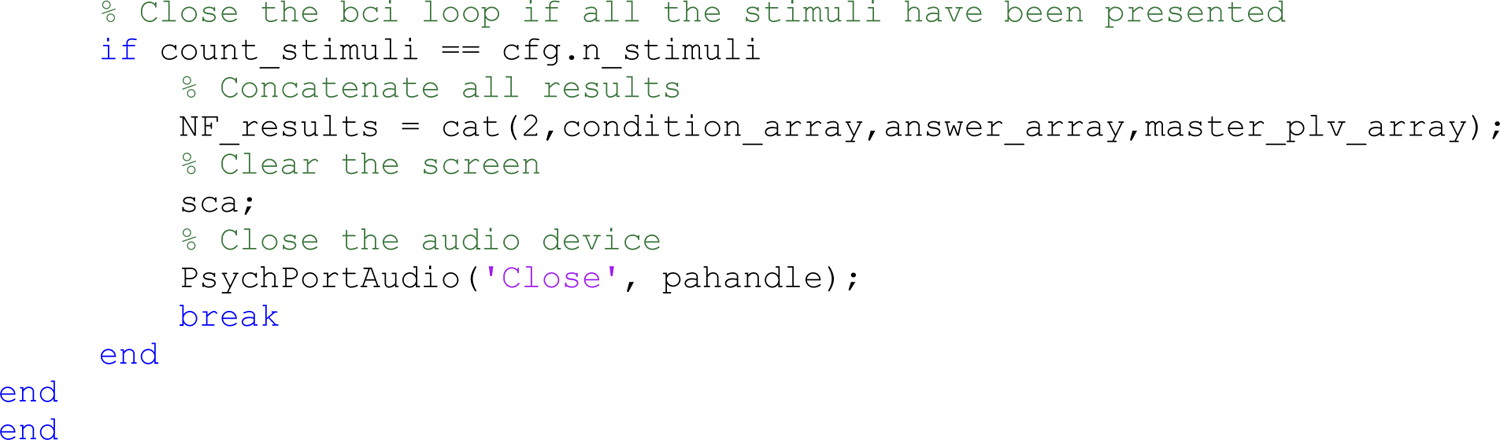

### Trigger_Gen_Multi.m

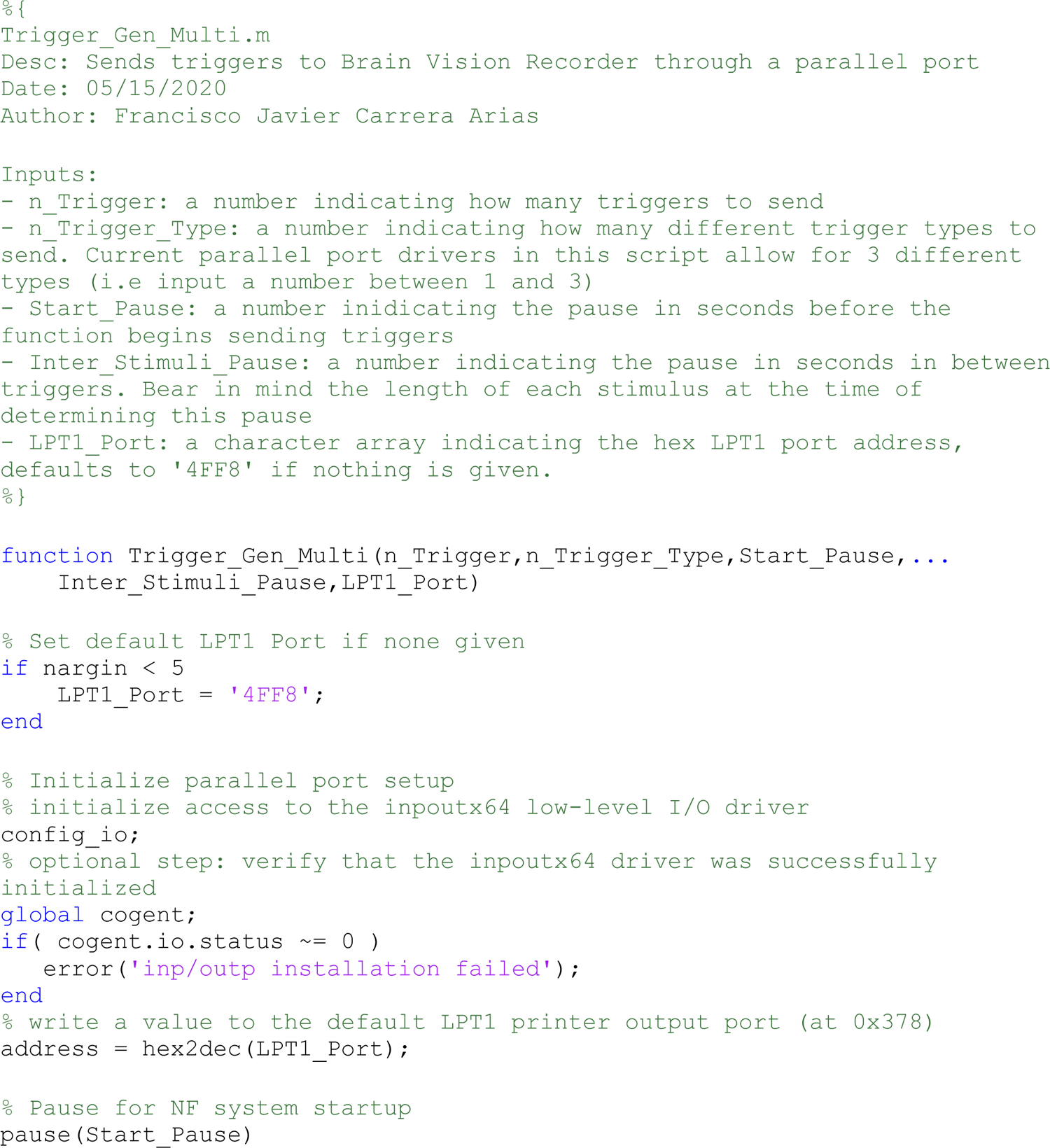
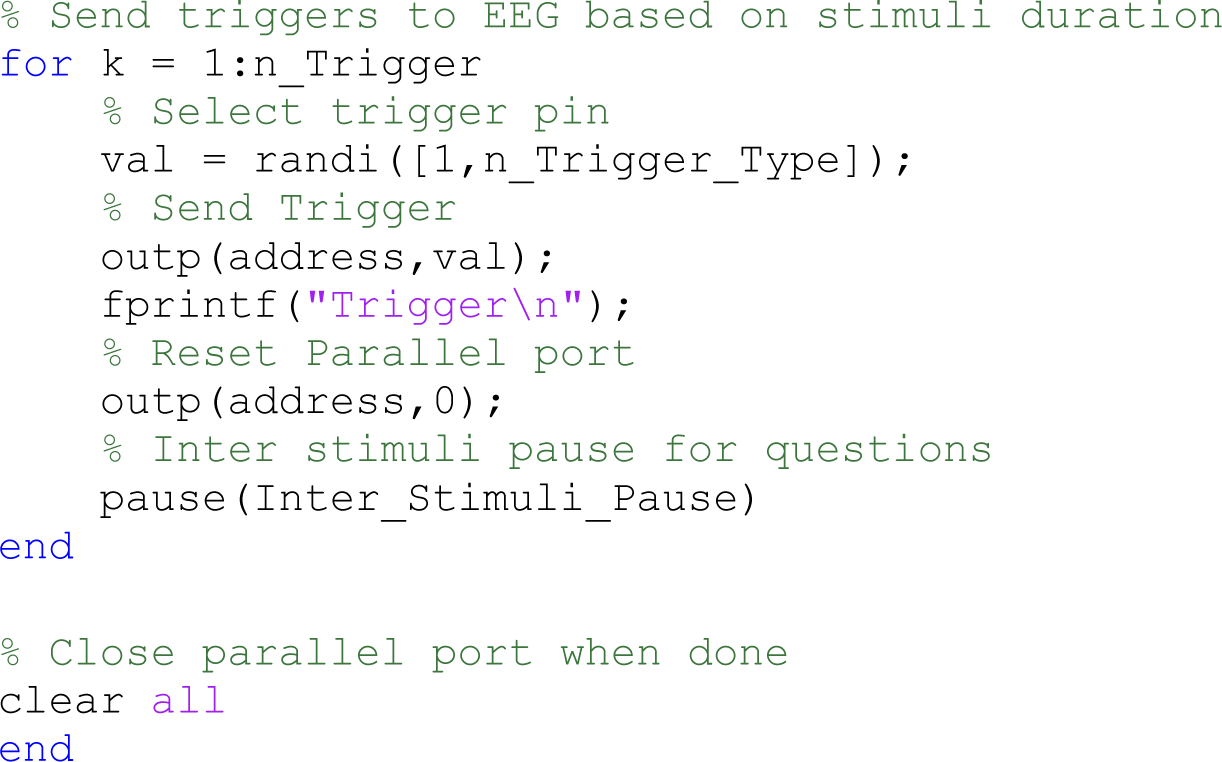

